# Estropausal gut microbiota transplant improves measures of ovarian function in adult mice

**DOI:** 10.1101/2024.05.03.592475

**Authors:** Minhoo Kim, Justin Wang, Steven E. Pilley, Ryan J. Lu, Alan Xu, Younggyun Kim, Minying Liu, Xueyan Fu, Sarah L. Booth, Peter J. Mullen, Bérénice A. Benayoun

## Abstract

Decline in ovarian function with age not only affects fertility but is also linked to a higher risk of age-related diseases in women (*e.g*. osteoporosis, dementia). Intriguingly, earlier menopause is linked to shorter lifespan; however, the underlying molecular mechanisms of ovarian aging are not well understood. Recent evidence suggests the gut microbiota may influence ovarian health. In this study, we characterized ovarian aging associated microbial profiles in mice and investigated the effect of the gut microbiome from young and estropausal female mice on ovarian health through fecal microbiota transplantation. We demonstrate that the ovarian transcriptome can be broadly remodeled after heterochronic microbiota transplantation, with a reduction in inflammation-related gene expression and trends consistent with transcriptional rejuvenation. Consistently, these mice exhibited enhanced ovarian health and increased fertility. Using metagenomics-based causal mediation analyses and serum untargeted metabolomics, we identified candidate microbial species and metabolites that may contribute to the observed effects of fecal microbiota transplantation. Our findings reveal a direct link between the gut microbiota and ovarian health.

## Main

Women are born with a definitive number of ovarian follicles and the gradual decrease in the number of the follicles ultimately leads to menopause^1–3^. The onset of menopause has been linked with an elevated risk of numerous age-related conditions, including osteoporosis and dementia^4–6^. Moreover, a later onset of menopause has been identified as a potential predictor of extended lifespan^7,8^. While the profound effects of menopause on the health- and lifespan of women are well-documented, the factors that contribute to ovarian aging and the strategies for its prevention remain poorly understood.

During the reproductive years, the hypothalamus-pituitary-ovarian hormone axis ensures the proper production and release of hormones necessary for ovulation and menstruation^9^. However, as menopause progresses, ovaries become less responsive due to reduced follicle numbers, ultimately leading to the hormonal imbalances characteristic of menopause^10^. Specifically, the serum concentrations of follicle-stimulating hormone (FSH) and luteinizing hormone (LH) increase, whereas anti-Müllerian hormone (AMH) and Inhibin A (INHBA) decrease with menopause progression^11–13^. These hormonal adjustments are hallmarks of the menopausal transition and highlight the diminished function of the ovaries^10^. Importantly, this disrupted feedback mechanism has been attributed to the wide array of menopausal symptoms and health risks associated with this transition phase^14,15^. For instance, recent studies have shown FSH receptor activation by FSH in the brain increases susceptibility for Alzheimer’s disease pathogenesis in women and blockade of FSH improves cognition in a mouse model of Alzheimer’s disease^14,16^.

The adult human gut microbiome is composed of approximately 10^13^ to 10^14^ micro-organisms^17,18^. The microbiome consists of bacteria, viruses and fungi, with bacteria representing the largest proportion of the microbial population^19^. Perturbations in the gut microbiota configuration are reported to lead to development and/or progression of serious disease, including diabetes and depression^20,21^. Interestingly, reported health effects of fecal microbiota derived from aged individuals have been inconsistent^22,23^. For example, one study demonstrated that fecal microbiota transplantation (FMT) from 24-month-old mice to 3-month-old mice led to the acceleration of age-associated phenotypes in the brain, retina, and gut of the recipient mice^22^. In contrast, another study found that grafting of fecal microbiota from 24-month-old mice to 5- to 6-week-old germ-free mice enhanced neurogenesis in the hippocampus and induced intestinal growth^23^. These findings underscore the microbiome’s complex, multifaceted effects on various organ systems throughout life, significantly affecting host health.

A recent study discovered that women with premature ovarian insufficiency exhibited significantly altered gut microbial profiles compared to healthy women^24^. Notably, alternations in the gut microbiome in premature ovarian insufficiency patients were associated with changes in serum hormone levels of key ovarian endocrine markers, including FSH, LH and AMH^24^. Similarly, the composition of the gut microbiome has been implicated in the development of polycystic ovary syndrome (PCOS), a disorder characterized by hyperandrogenism and polycystic ovaries^25^. Specifically, FMT of samples from PCOS patients to mice resulted in ovarian dysfunction in otherwise healthy recipient mice^25^. Consistent with the importance of a healthy microbiome for ovarian function, another study has shown that 17β-estradiol levels are significantly lowered in the germ-free female mice^26^. Moreover, transcriptome analysis of sexual development marker genes of the liver and histological studies of ovarian follicle development in germ-free female mice indicated that sexual maturation is perturbed in microbiota-depleted mice^26^. Importantly, the gut microbiota has been shown to modulate serum estrogen levels (through the “estrobolome”), further highlighting the potential role of the gut microbiota in female reproductive health^27^. The gut microbiota regulates action of estrogens through the secretion of β-glucuronidase, an enzyme that deconjugates estrogens into their active forms^27–29^. Dysbiosis of the gut microbiota can impair this process, leading to a reduction in circulating estrogens and potentially contributing to the development of estrogen-modulated conditions^27^. Collectively, these findings highlight a significant causal relationship between the gut microbiota and ovarian function and health, suggesting a pivotal role for microbial communities in modulating female reproductive conditions.

In this study, we analyzed the gut microbial profiles of young and estropausal (post-reproductive phase in rodents, analogous to human menopause) female mice via 16S rRNA amplicon sequencing and explored how the distinct microbial communities affect ovarian health through FMT experiments in young female mice. Unbiased RNA-seq analyses revealed that modifying the gut microbiota leads to profound changes in the ovarian transcriptomic landscape. Surprisingly, heterochronic gut microbiota transplant from estropausal mice to young female recipients led to a reduction in inflammation-related gene expression in the ovaries, and dampened expression of genes that tend to be upregulated in aging ovaries. Consistently, we observed improved ovarian function and health in the estropausal FMT recipients, supporting a surprising finding whereby old gut microbiota has beneficial effects on ovarian health. Lastly, we identified specific microbial genera and species, through 16S rRNA amplicon sequencing and whole genome shotgun sequencing, respectively, that might drive the observed transcriptomic changes in the ovaries via causal mediation analyses. Overall, our study shows that manipulating the gut microbiota can directly influence ovarian function and health.

## Results

### Young and estropausal female mice show distinct fecal microbial profiles

To investigate the potential link between changes in the gut microbiome and ovarian aging in mice, we assessed (1) the ovarian health state and (2) microbial profiles of young (YF, 4-months) and naturally aged estropausal (EF, 20-months) C57BL/6JNia female mice^30^ (Fig. 1A). To quantify the overall ovarian health state of the mice, we developed a composite “ovarian health index”, modeled after the concept of frailty index^31^ (Extended Data Fig. 1A). Our goal in constructing this index was to create a standardized, integrative tool that would enable comparative analysis of ovarian function across different datasets. The index combines ovarian follicle counts and serum levels of AMH, FSH, and Inhibin A, using a 3-tier scoring system based on the median values of the young and estropausal groups as reference points. Specifically, values beyond the young median are scored as 3, values between the medians of the young and estropausal groups are scored as 2, and values beyond the estropausal median are scored as 1 (Extended Data Fig. 1A). Given that ovarian follicle count and serum levels of key pituitary and ovarian hormones—AMH, FSH, and Inhibin A—are widely recognized as reliable indicators of ovarian health^32–35^, we combined these parameters to form a comprehensive and objective evaluation of ovarian function (Fig.1A-D; see Methods). Additionally, to facilitate accessibility and broader use of the ovarian health index, we developed an R shiny application, allowing the public to analyze their own datasets using this tool (https://alanxu-usc.shinyapps.io/Ovarian_Health_Index_Calculator/).

**Fig 1.**
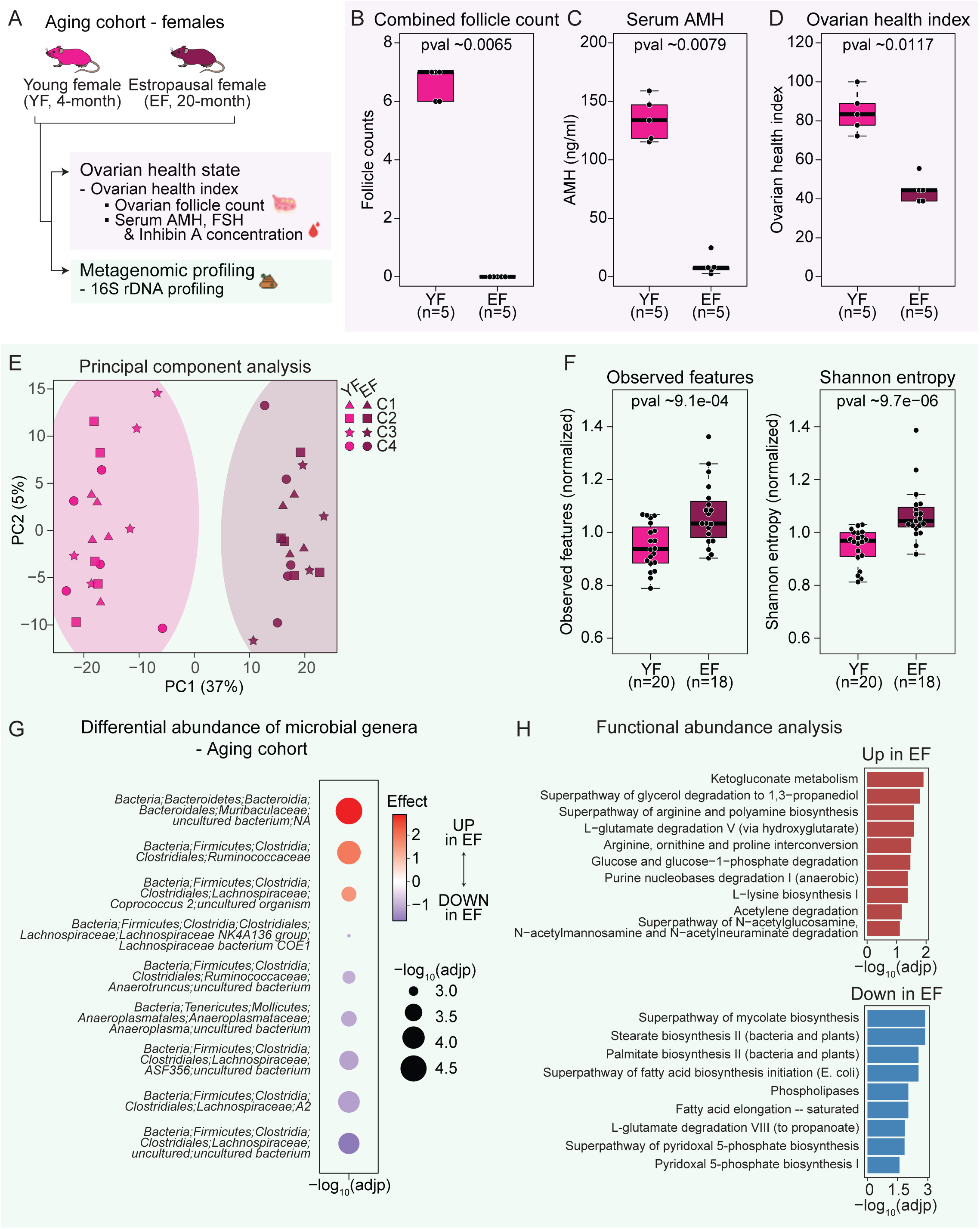
Characterization of ovarian health state and fecal microbial profiles of young and estropausal female mice. (A) Schematic diagram of the experimental setup of female aging cohorts. Resources from Flaticon.com were used. (B) Boxplot of combined follicle count of young and estropausal mice. (C) Boxplot of serum concentrations of AMH. (D) Boxplot of ovarian health index of young and estropausal mice. (E) Principal component analysis result of CLR-transformed and batch-corrected ASV counts from young and estropausal female mice. Animals from four independent cohorts (C1-C4), n=19-20 per group (variation due to animal death prior to experiment; n=20 for young females; n=19 for estropausal females). (F) Boxplots of Observed features and Shannon entropy indices of young and estropausal female mice. Medians from young females for each cohort were used to normalize indices per cohort. (G) Differential abundance analysis results of microbial genera of young vs. estropausal female mice using ALDEx2. (H) Functional abundance prediction analysis of young vs. estropausal female mice using PICRUSt2. Significance in nonparametric two-sided Wilcoxon rank-sum tests are reported for (B) and (D). YF: Young female; EF: Estropausal female. C: Cohort.

Consistent with previous reports^30,36^, we observed a significant reduction in ovarian follicle counts and serum levels of AMH and Inhibin A, as well as increased FSH levels in the estropausal group (Fig.1B,C and Extended Data Fig. 1B-D). Furthermore, the estropausal group exhibited a significantly lower ovarian health index score compared to the young group (p-value ∼0.0117; Fig.1D), validating the subsequent use of our index as a measure of ovarian health and function.

We then collected fecal samples from four independent cohorts of mice and performed 16S rRNA V3-V4 amplicon sequencing to compare the gut microbial profiles of the young and estropausal female mice. 16S rRNA amplicon sequencing allows identification of bacteria at the genus level by amplifying and sequencing variable regions of the 16S rRNA gene (*e.g*. V3-V4 region), which contains unique sequences specific to different species^37^. Quantification and analysis of the sequences from the variable regions, or amplicon sequence variants (ASVs), have been shown to provide important insights of the microbial compositions and environment^38^.

Our analysis focused on both beta diversity, which assesses differences between microbial communities, and alpha diversity, which looks at diversity within a community^39,40^. Beta diversity was evaluated using the Bray-Curtis dissimilarity and Jaccard index (Extended Data Fig. 2A). The analysis of Bray-Curtis dissimilarity and Jaccard index, conducted by cohort, revealed a distinct separation between the young and estropausal female groups (Extended Data Fig. 2A). Moreover, the results of the principal component analysis, applied to center log-ratio (CLR)-transformed and batch-corrected ASV counts, further highlighted clear clustering between the young and estropausal females across 4 independent cohorts of animals (Fig. 1E). Batch correction was performed to account for technical variability across cohorts^41^. Alpha diversity measures the diversity within a single microbial community or ecosystem, focusing on the variety and abundance of ASVs in a specific environment^36^. Importantly, estropausal females exhibited significantly higher scores in both observed features and Shannon entropy compared to the young female group (Fig. 1F). Our results suggest that the estropausal females have a higher biodiversity and increased species richness and evenness. This pattern of increased alpha diversity with age in female mice aligns with findings from previous studies^42^. However, in humans, alpha diversity has been shown variable trends with age, with both increases and decreases reported in different studies^43,44^. Additionally, a recent study reported elevated alpha diversity indices in PCOS model mice^45^.

We next asked which specific microbial genera changed in abundance between young and estropausal female mice recurrently across our 4 independent cohorts (Fig. 1G and Extended Data Fig. 2B, 3). Our analysis identified three genera that were more abundant (“UP in EF”) and six that were less abundant (“DOWN in EF”) in the microbiota of estropausal females (Fig. 1G and Extended Data Fig. 2B, 3; absolute effect size > 1, adjusted p-value < 0.05). Most of the differentially abundant genera belonged to the *Lachnospiraceae* family, offering a potential lead for future research on their impact on ovarian health. Previous studies have reported a relative decrease in abundance of microbial species from the *Lachnospiraceae* family with age in humans^46,47^. Further studies will be required to elucidate the molecular mechanisms through which these genera affect ovarian function and health. Additionally, we conducted a predictive analysis of functional consequences of these changes in microbial abundances using PICRUSt2^48^ (Fig. 1H). This analysis suggested an increase in metabolic pathways such as ketogluconate metabolism and the superpathway of glycerol degradation to 1,3-propanediol, while pathways including the superpathway of fatty acid biosynthesis initiation and phospholipases were predicted to decrease in the estropausal group (Fig. 1H). These predicted metabolic changes, including an increase in ketoglutarate and alterations in lipid metabolism, have been previously associated with ovarian cancers, underscoring the potential relevance of our findings to understanding ovarian health^49,50^. Our results collectively underscore the pronounced differences in the fecal microbial profiles between young and estropausal females.

### Chemically-induced premature ovarian failure model mice show distinct fecal microbial profiles

The 4-VinylCyclohexene Diepoxide (VCD)-injected mouse model is a follicle-deplete, ovary-intact animal model that is known to recapitulate many key aspects of human ovarian aging^51,52^. VCD accelerates atresia by causing selective loss of primary and primordial follicles in the ovary, while leaving the rest of the ovarian structure intact for residual hormone production^51,52^. We assessed the ovarian health index and fecal microbial profiles of vehicle- and VCD-exposed mice (CTL and VCD, respectively; Extended Data Fig. 4A). Briefly, 4-month-old C57BL/6NTac female and male mice were subjected to daily intraperitoneal injections of either vehicle (safflower oil) or VCD (at a dosage of 160 mg/kg/day) for 15 consecutive days. Then, the mice were maintained for an additional 100 days to ensure the complete development of phenotypes associated with VCD exposure^53^ (Extended Data Fig. 4A). To determine whether the observed effects on microbial profiles in females were a direct consequence of VCD-induced ovarian dysfunction rather than off-target effects of VCD administration, we also generated and analyzed data from male mice.

For female mice, ovarian follicle depletion and alterations in crucial ovarian hormone levels (*i.e.* AMH, FSH, and Inhibin A) were confirmed through ovarian histology analysis and serum hormone concentration measurements, respectively (Extended Data Fig. 4B-E). In line with these findings, the VCD-treated group exhibited a markedly lower ovarian health index (p-value ∼0.0104; Extended Data Fig. 4F), consistent with overall depleted ovarian function.

Subsequently, we conducted 16S rRNA V3-V4 amplicon sequencing on vehicle- and VCD-treated animals. Analysis through principal component analysis of CLR-transformed ASV counts, along with Bray-Curtis dissimilarity and Jaccard index, demonstrated a distinct separation between the vehicle-treated (CTL) and VCD-exposed groups (Extended Data Fig. 5A-B). Alpha diversity indices, on the other hand, revealed no significant differences between these two groups (Extended Data Fig. 5C). These findings suggest that the overall diversity within the group’s communities may appear similar, but they may be different in the species present, their relative abundances and the functions the species perform. Notably, beta diversity analysis of microbial profiles in male mice showed no notable distinctions between vehicle- and VCD-treated groups (Extended Data Fig. 5A, B). Additionally, there were no differences in alpha diversity between these groups (Extended Data Fig. 5D). These findings indicate that the observed effects on microbial profiles in females are causally induced by ovarian dysfunction and not off-target effects of VCD administration.

Our analysis revealed differentially abundant genera from CTL vs. VCD groups (Extended Data Fig. 5E and 6; absolute effect size > 1, adjusted p-value < 0.05). We detected several genera from the *Lachnospiraceae* family, yet none of the unique ASV IDs corresponded with those identified as differentially abundant in the estropausal female mice. Thus, further research to accurately identify the species significant to ovarian aging and health will be important. Then, we utilized PICRUSt2 to predict functional abundance between the CTL and VCD groups (Extended Data Fig. 5F). Notably, we detected down-regulation of the phospholipases pathway in the VCD group, a finding consistent with observations in the estropausal female group (Fig. 1H and Extended Data Fig. 5F). Our findings suggest that physiological ovarian aging and chemically-induced ovarian failure lead to substantially different outcomes on the gut microbiota.

### Young and old male mice show distinct fecal microbial profiles

Next, as an important comparison point, we also characterized the fecal microbial profiles of young (4-months) and naturally aged old (20-months) C57BL/JNia male mice (Extended Data Fig. 7A). This comparative study among female and male aging mice, as well as the VCD-exposed mice, aimed to identify differences in microbial profiles attributable to general aging and those specific to ovarian aging. We collected fecal samples from three independent cohorts of mice and performed 16S rRNA V3-V4 amplicon sequencing to compare the gut microbial profiles of the young and old male mice. Analysis using principal component analysis of CLR-transformed ASV counts, coupled with Bray-Curtis dissimilarity and Jaccard index, showed a distinct divergence between the young and old male groups (Extended Data Fig. 7B-C). However, this separation was less pronounced compared to what was observed in the female cohorts (Fig. 1E). Alpha diversity indices indicated no significant disparities between the two age groups of males (Extended Data Fig. 7D). Similarly, a previous study examining microbial profiles in male mice aged 3-4 months and 19-20 months found comparable alpha diversity index values between the two age groups^54^. Through differential abundance and functional abundance prediction analyses, we identified several genera and functional terms that varied significantly between young and old male mice (Extended Data Fig. 7E-F and 8). Importantly, we observed a few consistent functional terms across female and male aging cohorts. In particular, the L-lysine biosynthesis I pathway was found to be up-regulated, while the pyridoxal 5-phosphate biosynthesis I pathway and the superpathway of pyridoxal 5-phosphate biosynthesis were down-regulated with age in both sexes (Fig. 1H and Extended Data Fig. 7F). These results suggest that these pathways may play roles in the general aging processes, rather than being uniquely related to ovarian aging. Further research will be necessary to elucidate the influence of these functional pathways on the aging process.

### Gut microbiota manipulation can induce changes in the ovarian transcriptome

To investigate the potential effects of gut microbiota on ovarian health, we conducted FMT experiments to directly assess how alterations in the gut microbiota after estropause may influence ovarian function. Specifically, young adult female mice (4-months) were subjected to treatment with anti-biotics/anti-mycotic (Ab/Am) treatment to eliminate their initial microbial environment, and then received fecal microbiota grafts from either young (FMT-YF) or estropausal (FMT-EF) female mice (Fig. 2A). Subsequently, the ovaries were harvested for bulk RNA-seq analysis (Fig. 2A). Based on previous research that has shown adverse effects of gut microbiota from donors with ovarian dysfunction, such as PCOS^25^, our initial hypothesis was that transplanting fecal matter from estropausal mice could potentially induce estropause-like conditions in the recipient mice.

**Fig 2.**
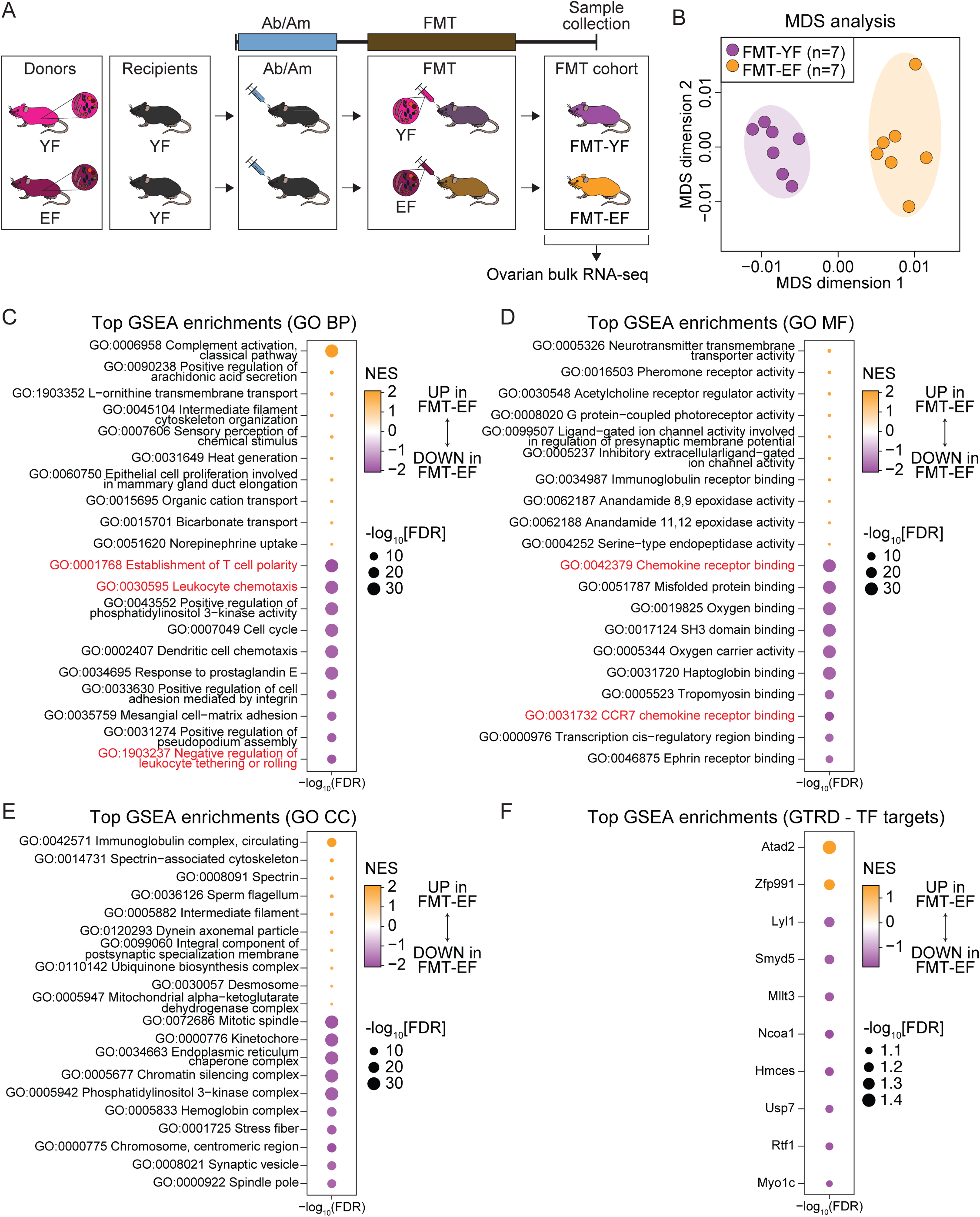
Bulk RNA-seq analysis of ovaries from young and estropausal female mice fecal microbiota transplantation recipient mice. (A) Schematic diagram of the experimental setup of fecal microbiota transplantation (FMT) cohort. Fecal microbiota from either young (4 months old) or estropausal (20 months old) female mice were transplanted into young (4 months old) female recipient mice. FMT recipient groups are referred to as FMT-YF (receiving young donor microbiota) and FMT-EF (receiving estropausal donor microbiota). (B) Multidimensional scaling (MDS) analysis result of RNA expression between young and estropausal female mice FMT recipient mice (FMT-YF and FMT-EF, respectively). (C-F) Top GSEA enriched terms from (C) Gene Ontology Biological Process (GO BP), (D) Gene Ontology Molecular Function (GO MF), (E) Gene Ontology Cellular Component (GO CC) and (F) Gene Transcription Regulation Database (GTRD – TF targets). Only up to top ten most-upregulated and top ten most-downregulated gene sets are plotted for readability.

Multi-dimensional scaling (MDS) analysis of the ovarian RNA-seq dataset revealed a clear separation between the FMT-YF and FMT-EF groups (Fig. 2B). Additionally, we detected 2,131 differentially expressed genes between the FMT-YF and FMT-EF groups (FDR < 5%; Extended Data Fig. 9A). Then, we investigated the pathways that were distinctively regulated between the FMT-YF and FMT-EF groups. Our gene set enrichment analysis (GSEA)^55^ results highlighted that pathways related to immune regulation were significantly down-regulated in the FMT-EF groups (Fig. 2C-E; highlighted in red). To evaluate whether these changes were the results of substantial changes in immune cells infiltration in the tissue, we leveraged a publicly available ovarian single-cell RNA-seq dataset^56^ to conduct deconvolution analysis, thus allowing us to deduce alterations in immune cell proportions within our dataset. Using two distinct tools, CSCDRNA^57^ and Granulator^58^, we determined that the proportions of immune cells did not significantly differ between the FMT-YF and FMT-EF groups (Extended Data Fig. 9B), consistent with the notion that the transcriptional changes that we observed may reflect cell autonomous changes in gene expression profiles. In addition, our GSEA results revealed terms associated with chromatin structure and dynamics, including kinetochore, chromatin silencing complex, chromosome, centromeric region, and spindle pole (Fig. 2E). These findings suggest that transcriptional changes in the ovaries may reflect alterations in chromatin organization and chromosome dynamics, processes critical for maintaining oocyte quality^59,60^. Together, these results indicate that pathways associated with both immune regulation and chromatin dynamics distinguish the FMT-YF and FMT-EF groups.

To identify transcription factors potentially driving the transcriptomic alterations observed between the FMT-YF and FMT-EF groups, we utilized Gene Transcription Regulation Database (GTRD)^61^ with the GSEA paradigm. While our analysis did not identify any transcription factors meeting the usual stringent criteria of FDR < 5%, we found two transcription factors showing trends of up-regulation and eight displaying down-regulation trends from the FMT-EF group, with an adjusted FDR < 10% threshold (Fig. 2F). Then, we identified publicly available ChIP-seq datasets for the trending transcription factors via the Cistrome Data Browser^62,63^, using corresponding *bona fide* peaks to determine potential enrichment by GSEA. Our analysis showed significant negative enrichment scores of Ncoa1 and Usp7-regulated genes within the FMT-EF group (p-value ∼0.0138 and ∼1e-04, respectively; Extended Data Fig. 9C). Moreover, we noted that the expression levels of *Ncoa1* and *Usp7* remained consistent across FMT-YF and FMT-EF groups (Extended Data Fig. 9D), suggesting potential post-transcriptional or post-translational regulation. While research on the roles of Ncoa1 and Usp7 in ovarian function is sparse, one study highlighted that Ncoa1 function is important in egg production traits in Shaobo hens^64^. Additionally, an inhibitor of Usp7, P5091, was found to inhibit the proliferation of ovarian cancer cells in another study^65^. Together with our findings, this suggests that Ncoa1 and Usp7 represent potential transcriptional regulators of ovarian function.

### Heterochronic FMT may have rejuvenating effects on the ovarian transcriptome of the recipients

Cumulative studies have indicated that aging leads to a pro-inflammatory shift in the ovarian microenvironment, adversely affecting ovarian function and oocyte quality^66,67^. Based on our GSEA results which indicated down-regulation of inflammatory pathways in the ovaries of the FMT-EF group, we next asked whether changes in the ovarian transcriptome upon FMT-EF could represent a more general trend of transcriptional rejuvenation. For this purpose, we analyzed two publicly available bulk RNA-seq datasets profiling ovaries from young (2-3-months) and aged/estropausal (12-months) wild-type female mice^68,69^ to identify gene sets that are either up- or down-regulated with ovarian aging (Fig. 3A). To evaluate whether the genes identified from publicly available RNA-seq datasets on ovarian aging (“UP with ovarian aging” and “DOWN with ovarian aging”) were significantly regulated in response to FMT-EF, we conducted GSEA using significant age-regulated genes as input gene sets. Surprisingly (and contrary to expectations based on previously reported observation at the somatic level for heterochronic FMT), our findings revealed a significant negative enrichment score for genes that were up-regulated in the context of ovarian aging (p-value ∼0.0046; Fig. 3B), suggesting that the ovarian transcriptome of the heterochronic FMT recipients (*i.e.* FMT-EF group) displays transcriptional profiles resembling those of younger females.

**Fig 3.**
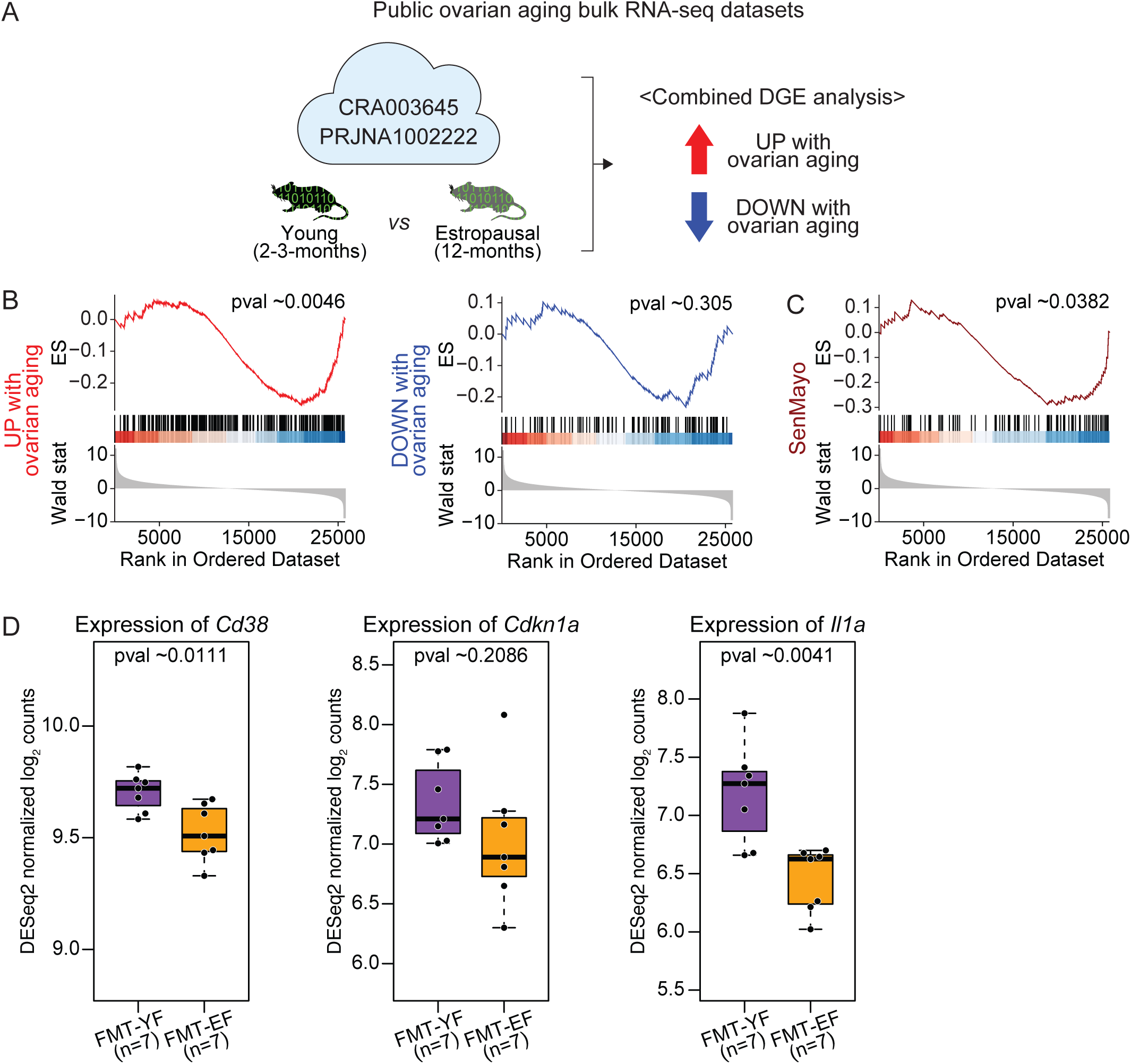
Analysis of public ovarian aging bulk RNA-seq datasets. (A) Schematic diagram of the analysis setup of public ovarian aging bulk RNA-seq datasets, CRA003645 and PRJNA1002222, from young (2-3-months) and estropausal (12-months) female mice. (B-C) GSEA enrichment score plots of (B) genes that are up-regulated with ovarian aging (“UP with ovarian aging”, left panel) and down-regulated with ovarian aging (“DOWN with ovarian aging, right panel), and (C) SenMayo gene set. In both (B) and (C), the black vertical lines indicate the positions of the genes from the gene set within the ranked list of genes. The curve represents the running enrichment score (ES), with the peak indicating the point of maximum enrichment for the gene set. The color scale at the bottom of the plot reflects the ranking metric, where red represents upregulated genes and blue represents downregulated genes. The p-value shown is calculated based on GSEA and adjusted for multiple testing using the Benjamini-Hochberg method. DGE: Differential gene expression. (D) Boxplots of DESeq2 normalized log_2_ counts of *Cd38*, *Cdkn1a* and *Il1a*.

As an orthogonal line of evidence, we next asked whether indicators of cellular senescence (as measured by the highly curated senescent marker gene set, SenMayo^70^) were significantly impacted by FMT-EF. Importantly, our results showed a significant negative enrichment score for the SenMayo gene set in the FMT-EF group, consistent with an overall “rejuvenation” trend for the ovarian transcriptome of FMT-EF mice (p-value ∼0.0382; Fig. 3C).

A recently published study highlighted the role of Nicotinamide Adenine Dinucleotide (NAD+) metabolism in ovarian aging, showing that increased NADase CD38 expression accelerated ovarian aging ^69^. CD38 deletion or pharmacological inhibition preserved fertility and follicle reserves in aged mice ^69^. Building on these findings, we examined the expression of *Cd38* as well as inflammation markers discussed in the study, including *Cdkn1a* and *Il1a* (Fig. 3D). In our analysis, we observed reduced *Cd38* expression in the FMT-EF group compared to the FMT-YF group, along with decreased expression of the inflammation markers (Fig. 3D). These findings suggest the potential involvement of the CD38-NAD pathway in the observed phenotypes of the FMT-EF group.

### FMT-EF mice show improved ovarian health

Since we observed large ovarian transcriptome remodeling in the FMT-EF group consistent with a less inflammatory, more youthful ovarian state, we next assessed the ovarian health state of the FMT-YF *vs.* FMT-EF groups to determine whether FMT-EF led to enhanced ovarian function (Fig. 4). We evaluated ovarian health function using a 2-pronged approach (Fig. 4A): (i) calculating the ovarian health index, and (ii) evaluating effective fertility using litter size and time to first pregnancy after mating^71^ (Fig. 4).

**Fig 4.**
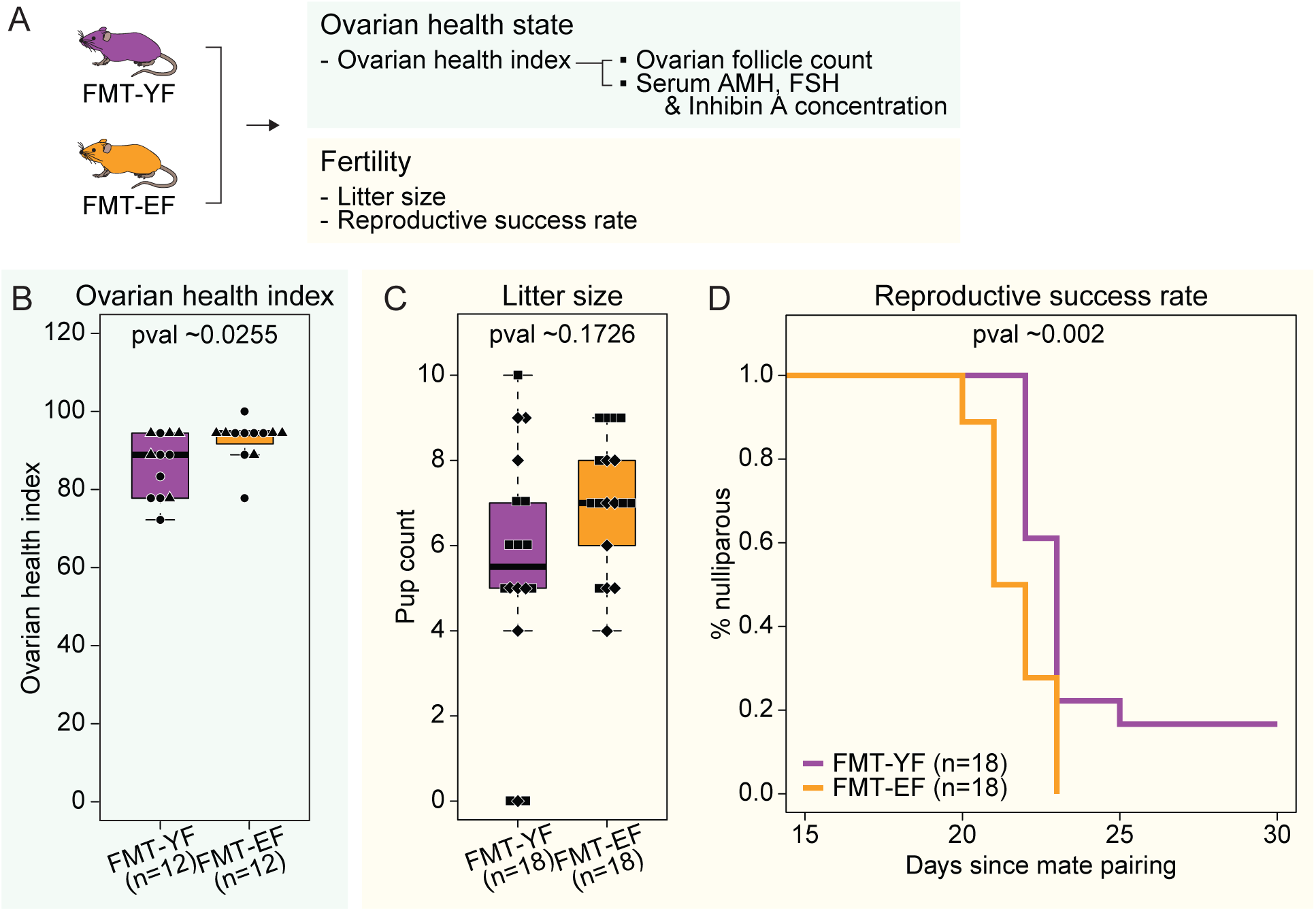
Evaluation of fertility and ovarian health state of fecal microbiota transplantation recipient mice. (A) Schematic diagram of the experimental setup. (B) Boxplot of ovarian health index of FMT-YF and FMT-EF mice. Ovarian health index was calculated for two independent cohorts (n=5 or 7 per group and 10 or 14 per cohort). (C) Boxplot of pup counts of first litters of young and estropausal female mice fecal microbiota transplantation recipient mice (FMT-YF and FMT-EF, respectively). Pup counts were measured from two independent cohorts (n=8 or 10 per group and 16 or 20 per cohort). (D) Reproductive success rate analysis result shown as percentages of nulliparous between FMT-YF and FMT-EF. Significance in nonparametric two-sided Wilcoxon rank-sum tests are reported for (B) and (C), and logrank test for (D).

Consistent with our prediction from the ovarian transcriptomes, the FMT-EF mice exhibited significantly higher scores on the composite ovarian health index than FMT-YF mice (p-value ∼0.0255; Fig. 4B, Extended Data Fig. 10A-D), supporting an enhanced ovarian health phenotype. The ovarian health index incorporates multiple parameters, including serum levels of AMH, FSH, and Inhibin A, as well as ovarian follicle counts (Extended Data Fig. 1A). Notably, for the FMT cohorts, FSH levels were measured using two different assay kits during the study due to an update implemented by the core facility conducting the assays. To ensure consistency and comparability of data across cohorts, we applied a correction (see Methods). This adjustment ensured that FSH levels accurately reflected ovarian health and were comparable between cohorts (Extended Data Fig. 10D). Next, we evaluated the result of ovarian function – fertility – through measures of litter size and reproductive success rate. Specifically, there was a suggestive trend for increased litter size in FMT-EF mice (p-value ∼0.1726; Fig. 4C). In addition, FMT-EF mice remained nulliparous for a shorter amount of time after pairing with a young health male (p-value ∼0.002; Fig. 4D), also suggestive of improved fertility. To determine whether the inclusion of 0 pup count data influenced our findings, we re-analyzed the latency data after excluding animals with 0 pup counts. Importantly, the latency data remained statistically significant (p-value ∼0.0159 without 0 pup counts vs. p-value ∼0.002 with all data). Together, our findings are consistent with improved/rejuvenated ovarian function after FMT-EF.

Importantly, no significant differences were observed in body weight between FMT-YF and FMT-EF groups after Ab/Am treatment (Extended Data Fig. 10E and Extended Data Table 1). Additionally, staging of the estrous cycle on the day of sample collection revealed no significant differences between the groups (Extended Data Table 2).

### FMT-YF and FMT-EF mice show distinct microbial profiles

Importantly, we next verified that fecal transplants from young or estropausal female mice led to distinct microbial profiles in the recipient mice. For this purpose, we conducted and analyzed 16S rRNA V3-V4 amplicon sequencing on the gut microbiota of FMT-YF and FMT-EF mice (Extended Data Fig. 11). Principal component analysis of CLR-transformed ASV counts, complemented by Bray-Curtis dissimilarity and Jaccard index, showed a clear segregation between the FMT-YF and FMT-EF groups (Extended Data Fig. 11B). While no significant differences were noted across four alpha diversity indices (Extended Data Fig. 11C), we identified several microbial genera with significant differential abundance between the FMT-YF and FMT-EF groups (absolute effect size > 1, adjusted p-value < 0.05; Extended Data Fig. 11D and 12). Notably, two genera that showed increased abundance in the FMT-EF recipient group were also elevated in the donor estropausal females (*Bacteria;Firmicutes;Clostridia;Clostridiales;Ruminococcaceae* and *Bacteria;Firmicutes;Clostridia; Clostridiales;Lachnospiraceae;Coprococcus 2;uncultured organism;* Extended Data Fig. 11E). These results indicate that transplanting microbiota from young or estropausal female mice led to changes in the microbial landscape of the recipient mice. Other studies have also reported the selective enrichment of specific microbial species from donor microbiota in recipient subjects^22,72^. These observations may be attributable to variations in the intestinal microenvironments, which allow the proliferation of certain species.

Subsequently, we used PICRUSt2 to predict functional abundance differences between the FMT-YF and FMT-EF groups (Extended Data Fig. 11F). We identified significant functional groups only with up-regulated functional terms in the FMT-EF group, with no significant down-regulated terms. Notably, there was a significant presence of terms related to the menaquinol biosynthesis pathway, including those involved in 1,4-dihydroxy-6-naphthoate biosynthesis II, 1,4-dihydroxy-6-naphthoate biosynthesis I, and the superpathway of menaquinol-8 biosynthesis II (Extended Data Fig. 11F). Menaquinols, the active hydroquinone forms of vitamin K, also known as menaquinones (MKs), facilitate the carboxylation of vitamin K-dependent proteins, which are crucial for biological processes such as blood clotting, bone health, and heart health^73^.

To pinpoint specific microbial species that may contribute to the observed changing in the ovarian health state, we performed whole genome shotgun (WGS) metagenomics on the FMT cohort (Fig. 5A). Consistent with the 16S amplicon sequencing results, PCA analysis of CLR-transformed data and beta diversity analysis revealed distinct clustering between the FMT-YF and FMT-EF groups (Fig. 5B-C). Similarly, no significant differences were observed in alpha diversity indices, reinforcing the similarity in overall microbial diversity between the groups (Fig. 5D). WGS analysis identified 168 differentially abundant microbial species (FDR < 5%; Fig. 5E-F and Extended Data Fig. 13), further confirming distinct microbial compositions in the FMT-YF and FMT-EF groups. Intriguingly, microbial species previously reported to be differentially enriched in the fecal microbiota of PCOS patients, such as *Bacteroides xylanisolvens*, *Bacteroides thetaiotaomicron*, *Parabacteroides distasonis*, and *Bacteroides stercoris*^25^, were significantly depleted in the FMT-EF group (Extended Data Table 3).

**Fig 5.**
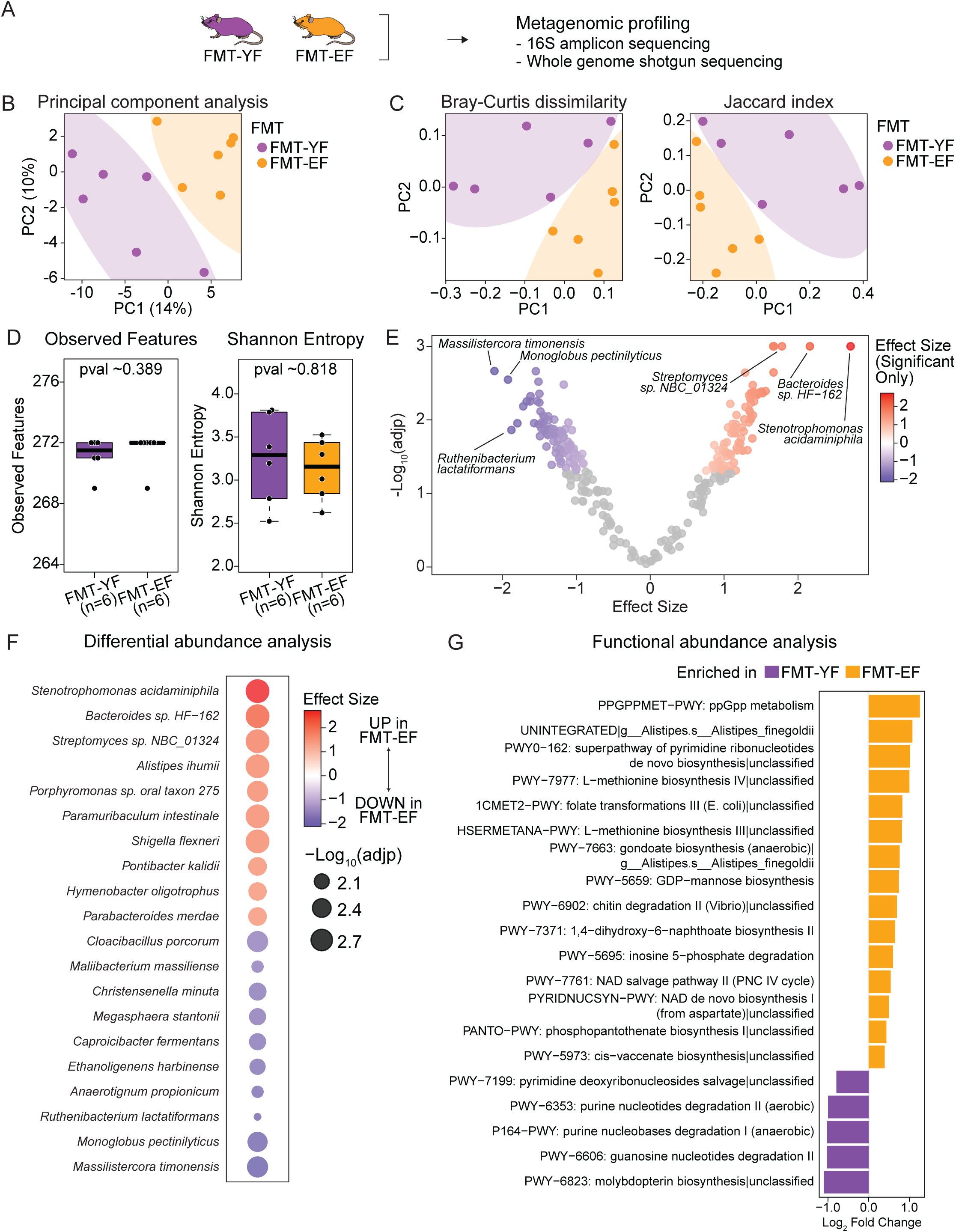
Characterization of fecal microbial profiles of fecal microbiota transplantation recipient mice. (A) Schematic diagram of the experimental setup. (B) Principal component analysis result of CLR-transformed and batch-corrected counts from young and estropausal female mice fecal microbiota transplantation recipient mice (FMT-YF and FMT-EF, respectively). (C) Principal coordinate analysis results of Bray-Curtis dissimilarity (left panel) and Jaccard (right panel) indices. (D) Boxplots of Observed features and Shannon entropy indices of FMT-YF and FMT-EF mice. (E) Volcano plot of differential abundance analysis results of microbial species of FMT-YF vs. FMT-EF mice using ALDEx2. (F) Bubble plot of top 10 up- and down-regulated microbial species in FMT-EF. (G) Functional abundance analysis of FMT-YF vs. FMT-EF mice using HUMAnN3 (Q-value < 0.1). Significance in nonparametric two-sided Wilcoxon rank-sum tests are reported for (D).

Functional abundance analysis using HUMAnN3^74^ revealed up-regulation of pathways related to menaquinol biosynthesis, including PWY-7371: 1,4-dihydroxy-6-naphthoate biosynthesis II, consistent with predictions from the 16S amplicon analysis (Fig. 5G and Extended Data Fig. 11F). Intriguingly, we detected terms associated with the NAD biosynthesis pathway, such as PWY−7761: NAD salvage pathway II (PNC IV cycle) and PYRIDNUCSYN−PWY: NAD de novo biosynthesis I (from aspartate)|unclassified, which aligns with our ovarian RNA-seq data showing altered expression of *Cd38* in the FMT-EF ovaries (Fig. 3D, 5G).

Given the stability of menaquinones and their primary synthesis by gut microbiota, we further evaluated their levels in stool samples (MK6-MK13) to explore potential systemic effects post-absorption, as they contribute significantly to the host’s vitamin K status. However, no significant differences in MKs concentrations in the stool samples of FMT-YF and FMT-EF animals were observed (Extended Data Fig. 14A,B), although this may be the result of high animal-to-animal variability and/or the high vitamin K content of the standard mouse chow at the USC vivarium dietary (see Discussion).

### FMT-YF and FMT-EF mice show distinct serum metabolomic profiles

It is well established that gut microbiota-derived molecules can enter the bloodstream and influence health^75^. Additionally, metabolite levels in plasma have been shown to be affected by aging^76^. To assess whether shifts in the gut microbial profiles of the FMT animals induced changes in circulating metabolites, we performed untargeted metabolomics on serum samples from two independent experimental cohorts (Fig. 6A). First, we performed PCA of the identified metabolites to detect unbiased separation between the FMT-YF and FMT-EF groups. Although non-linear batch effects were present and could not be fully corrected, a clear separation was still evident within each batch (Fig. 6B). Subsequent LDA analysis further showed the distinct metabolomic profiles of the two groups (Fig. 6C). Finally, we extracted metabolites that changed consistently across both cohorts and selected the top 15 based on the LDA loadings (Fig. 6D).

**Fig 6.**
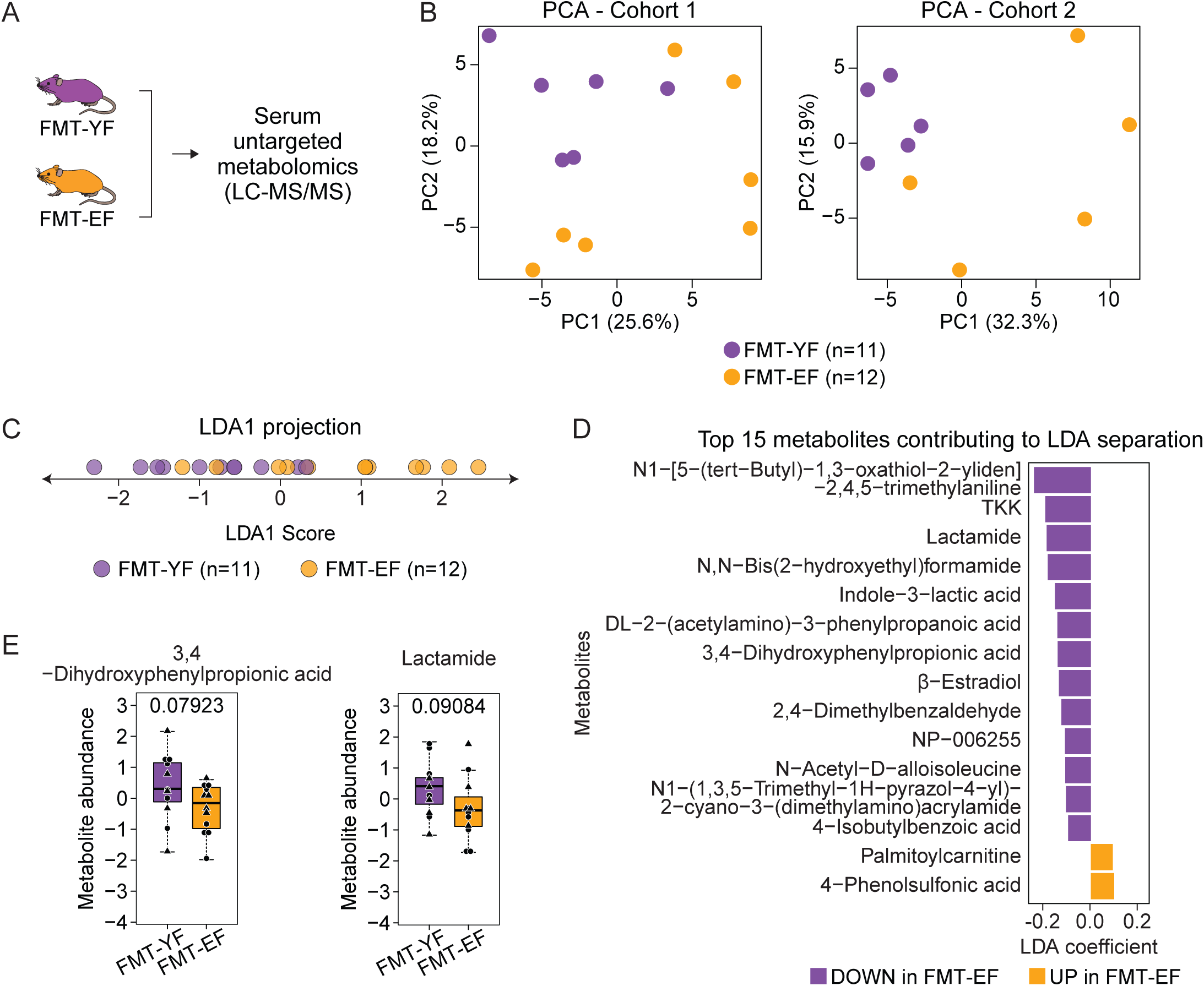
Characterization of serum metabolic profiles of fecal microbiota transplantation recipient mice. (A) Schematic diagram of the experimental setup. (B) PCA plots of identified serum metabolites from two independent FMT cohorts. (C) LDA2 projection scatter plot displaying LDA1 scores for the FMT-YF and FMT-EF groups. (D) Top 15 metabolites contributing to LDA-based separation, ranked by their positive and negative contributions to the LDA axis. (E) Boxplots of 3,4-dihydroxyphenylpropionic acid (3,4-DHPPA) and lactamide serum levels between FMT-YF and FMT-EF groups. Significance in nonparametric two-sided Wilcoxon rank-sum tests are reported for (E).

Among the metabolites identified, 3,4-dihydroxyphenylpropionic acid (3,4-DHPPA) and lactamide were significantly decreased in FMT-EF animals at FDR < 0.1 (Fig. 6D, E). 3,4-DHPPA, a microbial metabolite derived from polyphenol metabolism, has been linked to lipid metabolism and broader metabolic regulation^77^. Given the critical role of lipid metabolism in hormonal balance and ovarian function, its reduction in FMT-EF animals may reflect metabolic adaptations that contribute to improved ovarian health. Additionally, while the direct role of lactamide in ovarian function is unclear, microbial-derived lactic acid and its derivatives have been implicated in gut microbiota-driven regulation of reproductive health^78^. For instance, lactic acid bacteria have been shown to influence sex hormone-related gut microbiota composition, leading to beneficial effects on ovarian physiology in preclinical models^78^. The observed decrease in lactamide levels may indicate broader microbiome-mediated metabolic changes that support ovarian health.

In addition to these significant metabolic changes, palmitoylcarnitine, a key metabolite involved in fatty acid β-oxidation and mitochondrial energy production^79^, was elevated in FMT-EF animals according to LDA loadings. This suggests enhanced mitochondrial efficiency and metabolic flexibility, which are essential for oocyte and follicular health^80^. Furthermore, palmitoylcarnitine has been implicated in immune modulation^81^, and its increased levels may indicate a metabolic shift supporting both energy homeostasis and an anti-inflammatory ovarian environment. This aligns with our bulk RNA-seq analysis, which revealed a downregulation of inflammatory pathways in the ovaries of FMT-EF animals (Fig. 2C-E).

Conversely, LDA loadings for β-estradiol suggested that serum levels are decreased in FMT-EF mice. Based on the improved ovarian function observed in FMT-EF mice, this may reflect a more physiologically regulated hormonal environment, rather than a pathological decline. Estradiol plays a critical role in ovarian function, lipid metabolism, and mitochondrial activity^82^, and its levels may be influenced by gut microbiota-driven metabolic pathways^27^. The gut microbiome is known to modulate steroid hormone metabolism^27^, and FMT from EF donors may have contributed to a systemic reprogramming of lipid and hormonal pathways that promote ovarian resilience.

Together, these findings suggest that FMT from estropausal donors induces distinct metabolic shifts that may contribute to the observed improvements in ovarian health. Further studies will be needed to determine the functional impact of these metabolites on ovarian physiology and their potential role in reproductive aging.

### Causal mediation analyses reveal microbial species potentially influencing ovarian transcriptome

Lastly, we performed causal mediation analyses to characterize microbial species that may directly influence the ovarian transcriptome remodeling that we observed between FMT-YF and FMT-EF mice (Fig. 7A). From the features exhibiting significant abundance variations between the groups, we identified four microbial species (*Bacteroides zhangwenhongii, Shigella flexneri, Bacteroides stercoris*, and *Bacteroides graminisolvens*) with a significant mediation effect on ovarian transcriptome changes (Fig. 7B). Interestingly, *Bacteroides stercoris* was previously highlighted as a key contributor to the distinct fecal microbiota profiles between healthy individuals and those with PCOS^25^. Further investigation into the functional roles of these identified species in modulating ovarian function will be essential in future studies. Similarly, we identified four ASV IDs (850e0d58c1aa35238538564a8010e0ff, e4f32cdc3407edb8423447421078fdae, 55ebdcf041fb6b8b4632993423011372, and d4840c45eb37593108d020fb6a63f8e2) with a significant mediation effect on ovarian transcriptome changes (Extended Data Fig. 16).

**Fig 7.**
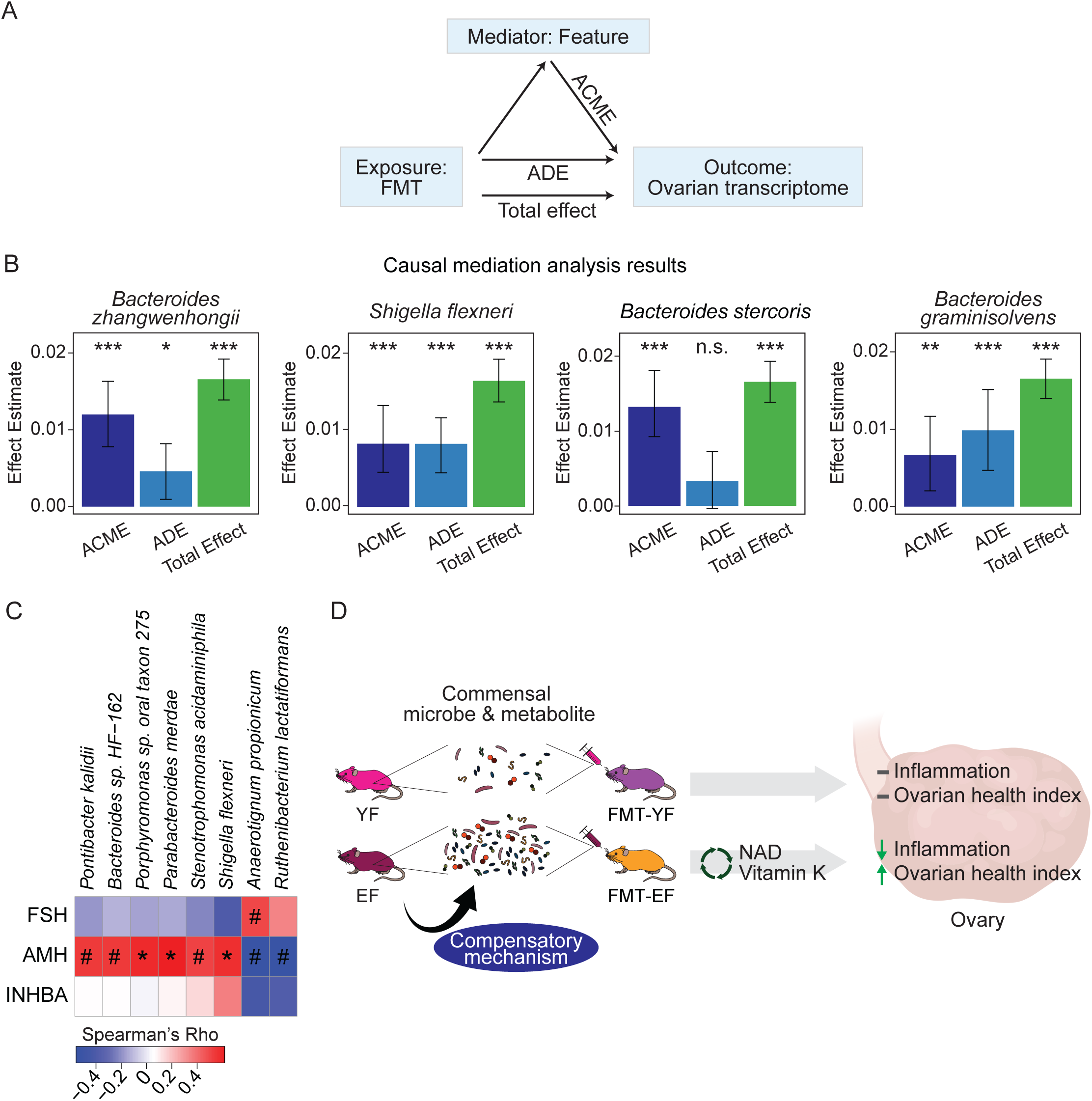
Causal mediation analysis of microbial species and ovarian transcriptome changes in fecal microbiota transplantation recipient mice. (A) Graphical representation of causal mediation analysis. (B) Causal mediation analysis results of whole genome shotgun metagenomics counts that showed significant mediation effects on changes in the ovarian transcriptome in fecal microbiota transplantation recipient mice. Nonparametric bootstrap confidence intervals with the percentile method are reported. (C) Heatmap of Spearman correlation analysis between CLR-transformed WGS count data and serum hormone quantification data. #: p-value < 0.1; *: p-value < 0.05. (D) Summary diagram. The estropausal female fecal microbiota exhibits increased levels of commensal microbes and metabolites through compensatory mechanisms. As a result, fecal transplantation of estropausal microbiota enhances ovarian health measures. Potential mechanisms, including NAD and vitamin K synthesis pathways, should be further examined. Generic Diagramming Platform (GDP) was used to generate the summary diagram^138^. ACME: Average Causal Mediated Effect; ADE: Average Direct Effect.

To further explore the relationship between gut microbiota and ovarian health, we performed Spearman correlation analysis using WGS metagenomics data and serum hormone quantification data (Fig. 7C). This analysis revealed significant correlations between multiple microbial species and hormone levels, including *Shigella flexneri, Parabacteroides merdae,* and *Porphyromonas sp. oral taxon 275,* suggesting these species as potential candidates influencing ovarian function. These findings provide a strong foundation for future studies aimed at experimentally validating the roles of these microbial species in modulating ovarian function and hormone levels.

## Discussion

In this study, we examined the fecal microbial profiles of three distinct groups: (1) young and naturally aged estropausal female mice, (2) mice with chemically-induced premature ovarian failure and their control counterparts, and (3) young and naturally aged male mice. Through our comparative analysis, we identified and differentiated between functional pathways that may be unique to ovarian aging (*i.e.* phospholipases pathway) and those specific to general aging (*i.e*. L-lysine biosynthesis I, pyridoxal 5-phosphate biosynthesis I, and superpathway of pyridoxal 5-phosphate biosynthesis). Furthermore, through fecal microbiota transplantation assays, we discovered that alterations in the gut microbiota can significantly impact the ovarian landscape, affecting both fertility and the health state of the ovaries. We also identified specific microbial species as potential mediators of the observed effects on ovarian function. The vitamin K2 and CD38-NAD pathways emerged as potential mechanisms to explore for investigating the underlying causes of the observed phenotypes in our FMT experiments (Fig. 7D). Future investigations focusing on these species hold significant promise for crafting novel therapeutic strategies aimed at mitigating or potentially reversing ovarian aging.

### Ovarian RNA-seq data of the FMT recipients supports causal influence of the gut microbiota on ovarian function

Our analysis of ovarian RNA-seq data from the FMT recipients (FMT-YF vs. FMT-EF) revealed significant changes in the ovarian transcriptome following alterations in the gut microbiota. While existing research underscores the broad effects of the gut microbiome on host health, evidence detailing its direct effects on the transcriptome of various tissues remains sparse. For instance, earlier research demonstrated that mice raised in a germ-free environment exhibit a markedly distinct liver transcriptome compared to their conventionally raised counterparts^26^. To the best of our understanding, the direct influence of gut microbiota on the ovarian transcriptome has not yet been documented. Our analysis not only uncovered significant variations in the ovarian transcriptome among FMT recipient mice (*i.e.* reduced inflammatory pathway in the FMT-EF group), but we also noted the transmission of phenotypic alterations (*i.e*. improved fertility and ovarian health). Collectively, these results offer substantial evidence of gut microbiota’s direct role in influencing ovarian function and health. Furthermore, the microbial species and genera, and functional pathways we identified serve as valuable assets for future research aimed at investigating potential therapeutic strategies.

### Unexpected advantageous effects of grafting estropausal female fecal microbiota may be due to compensatory mechanisms

Based on previous research that has shown detrimental effects of gut microbiota from donors with ovarian dysfunction, such as PCOS^25^, we initially hypothesized that transplanting fecal microbiota from estropausal mice might induce estropause-like phenotypes in the recipient mice. Contrary to our expectations, however, we observed an ovarian transcriptome in the recipients that resembled that of young ovaries, alongside improvements in ovarian health and fertility following the transplantation of microbiota from estropausal females. The unexpected advantageous effects observed could be attributed to a compensatory mechanism. For instance, FSH, required for follicle maturation and ovulation, is found in elevated levels in post-menopausal women and estropausal mice^2,83,84^. After reproductive senescence, the ovaries no longer produce estrogen or mature eggs, so the pituitary gland compensates by producing more FSH in an attempt to stimulate the now non-responsive ovaries^30^. Similarly, it is conceivable that the gut microbiota in reproductively senescent female mice could be enriched with beneficial microbes and/or metabolites that support ovarian health, perhaps as a compensatory response. Cumulative studies have highlighted the role of gut microbiota in compensatory processes. For instance, one study demonstrated that gut microbiota can influence the levels of peptides that regulate energy balance in the hypothalamus, serving as a mechanism to decrease body fat^85^. Furthermore, an increase in microbes that produce metabolites critical for the nervous system’s function—such as serotonin, acetylcholine, histamine, and dopamine—was observed in patients with mild cognitive impairment^86^. Therefore, when young adult female mice received fecal microbiota transplants from estropausal females, it is possible that the enriched gut microbial environment contributed to the manifestation of younger or more reproductively active phenotypes in recipient mice.

### Potential involvement of MKs in ovarian health and aging

Although our analysis of functional abundance from both WGS and 16S rRNA amplicon sequencing datasets predicted potential shifts in the biosynthesis pathway of MKs in the FMT mice, we observed no significant differences in the stool concentrations of MKs between the FMT-YF and FMT-EF groups. This lack of observable difference might be attributed to the vitamin K content of the standard chow used by the USC vivarium during our experiments (LabDiet, PicoLab Rodent Diet 20). Specifically, the high vitamin K dietary content could have saturated the system, thus masking subtle variations in MK production by the gut microbiome. Such nuances are typically more pronounced under conditions of a low vitamin K diet^87^. Published studies have shown that vitamin K status influences MK production because gut bacteria require vitamin K substrates to synthesize MKs^87^. In our study, the presence of vitamin K3 (menadione) in the diet, which is converted into MK4 in the body, likely contributed to the high basal MK levels observed in the fecal matter of the recipient mice. While there have been limited studies, MKs have demonstrated potential benefits for ovarian health in specific contexts, such as certain ovarian cancers and in patients with PCOS^88,89^. Additionally, some studies have underscored the anti-inflammatory properties of MKs^90^. Given our observations of decreased inflammatory function and enhanced ovarian function in the FMT-EF mice, coupled with the documented benefits of MKs, MKs may be therapeutic candidates for improving ovarian health and function. However, further research into the direct effects of MKs on ovarian function will be necessary.

### Sex-differences in aging-associated gut microbial profiles

In our comparative study, while we detected certain similarities in the microbial profiles of both females and males with aging, our analysis primarily revealed notable differences between the sexes. Specifically, while aging was associated with significant changes in alpha diversity in female mice, such changes were not observed in males. Moreover, our analyses of differential abundance of microbial genera and predictions of functional abundance revealed substantial differences in the gut microbial profiles and their functions between females and males as they age. Indeed, extensive research has established that aging manifests differently in females and males, a phenomenon known as sex dimorphism in aging^91^. Particularly, the susceptibility to diseases associated with aging, including Alzheimer’s disease and obesity, varies significantly between the sexes^92–94^. Furthermore, the disparity in lifespan between females and males is consistently observed across various species^95,96^. In agreement with our findings in this study, sex differences in the gut microbiome associated with aging have previously been documented^97,98^. While the limited number of existing studies suggest a possible influence of sex hormones on the observed sex differences in the gut microbial profiles^99–101^, these findings do not negate the potential effects of non-hormonal factors, such as those related to sex chromosomes, on the differences noted between sexes. Given the significant distinctions between sexes highlighted in this study and their varied impacts, it is imperative to consider sex as a contributing factor in future gut microbiome research, not only to deepen our understanding but also to address the specific differences effectively.

### The potential role of the CD38-NAD pathway in mediating ovarian rejuvenation in the FMT-EF mice

CD38, a primary NADase in mammalian tissues, plays a pivotal role in regulating NAD+ levels^102^. Emerging evidence implicates the CD38-NAD axis in aging, with increased CD38 expression contributing to age-related declines in NAD+ levels, impairing mitochondrial function and cellular metabolism^103,104^. In the context of ovarian aging, studies have shown that CD38 overexpression accelerates follicular depletion and diminishes ovarian function, whereas CD38 inhibition preserves follicle reserves and fertility in aged mice^69^.

In this study, we observed reduced *Cd38* expression in the ovaries of the FMT-EF group, alongside downregulation of inflammatory markers such as *Cdkn1a* and *Il1a* (Fig. 3D). Functional pathway predictions from the metagenomics data further supported these findings, revealing enrichment of pathways related to NAD biosynthesis in the FMT-EF group (Fig. 5G). Specifically, pathways such as NAD salvage pathway II (PNC IV cycle) and NAD de novo biosynthesis I (from aspartate) were upregulated (Fig. 5G). These results suggest that the gut microbiota established through estropausal fecal microbiota transplantation may suppress CD38 activity, influencing ovarian NAD metabolism and creating a more favorable ovarian environment. This aligns with previous research linking CD38-mediated NAD+ depletion to inflammation and cellular senescence, processes that are detrimental to ovarian function^105,106^.

Together, these findings indicate that modulation of the CD38-NAD pathway by gut microbiota may alleviate ovarian inflammation and promote a more youthful transcriptomic profile, contributing to the enhanced ovarian health observed in FMT-EF mice.

## Limitations of the study

One potential caveat in our chemically induced ovarian failure model is that VCD, while selectively depleting ovarian follicles, is a systemically administered agent and could potentially affect other biological systems. However, studies have shown that VCD’s effects are confined to ovarian tissue, and its impact on other organ systems have not been reported^107,108^. To further assess off-target effects, we conducted control experiments in male mice injected with VCD or vehicle, and no significant differences in gut microbial beta diversity indices (Bray-Curtis dissimilarity and Jaccard index) were observed. These findings suggest that the observed microbial changes in female mice are primarily due to ovarian dysfunction rather than direct effects of VCD, though we acknowledge the potential for non-specific effects. Additionally, it is possible that the improved fertility outcomes in the FMT-EF group, compared to the FMT-YF group, may reflect differences in how aged and young donor microbiota recolonize the gut following antibiotics treatment. For example, the aged microbiota might better adapt to recolonization, reducing the likelihood of artificially poor fertility outcomes in the recipients. Nonetheless, the changes in the microbiota induced by FMT resulted in significant changes in ovarian health, as evidenced by differences in ovarian transcriptomic profiles, fertility outcomes, and overall ovarian health indices in the FMT-YF and FMT-EF groups. Finally, another limitation of the study involves the use of PICRUSt2, which predicts functional abundances based on marker gene sequences. While PICRUSt2 provides insights into potential functional pathways affected by changes in the microbiota, it is a predictive tool and does not directly measure functional abundances. Future studies will require additional functional assays to validate these predictions.

In summary, we performed a comparative analysis of microbial profiles among aging female, aging male and a chemically-induced menopause model mice to characterize microbial genera that are unique to general aging vs. ovarian aging. Our work highlights potential microbial species and genera, and functional pathways that may play key roles in modulating ovarian function.

## Acknowledgements

We thank Dr. Victor Ansere and Dr. Juan Bravo for feedback on our manuscript. We also thank Dr. Dario Valenzano for his insightful comments and guidance on microbiome data analysis.

This work was supported by the GCRLE-2020 post-doctoral fellowship from the Global Consortium for Reproductive Longevity and Equality at the Buck Institute, made possible by the Bia-Echo Foundation to M.K., USC Provost’s Undergraduate Research Fellowship to J.W., National Institute on Aging (NIA) T32 AG052374 predoctoral fellowship and Diana Jacobs Kalman/AFAR Scholarships for Research in the Biology of Aging to R.J.L., and Pew Biomedical Scholar award #00034120 from the Pew Charitable Trust to B.A.B. Ovarian histological analysis was performed by the Translational Pathology Core at the USC Norris Comprehensive Cancer Center (supported by NCI P30 CA014089). Serum hormone quantification was performed by the University of Virginia Center for Research in Reproduction Ligand Assay and Analysis Core (supported by the Eunice Kennedy Shriver NICHD/NIH Grant R24HD102061). Computational analyses were performed using the Center for Advanced Research Computing (CARC) resources at the University of Southern California (USC). Stool vitamin K concentrations were supported by the USDA Agricultural Research Service under Cooperative Agreement No. 58-1950-7-707. Any opinions, findings, conclusions, or recommendations expressed in this publication are those of the authors and do not necessarily reflect the views of the USDA. We thank Whitaker Cohn, the Assistant Director of the USC Mann Multi-Omics Mass Spectrometry Core, for running the metabolomics samples on their instruments.

## Author contributions

M.K. and B.A.B. designed the study. M.K. performed live animal experiments. M.K. and R.L.J. dissected mouse tissues. M.K., J.W. and Y.K analyzed ovarian histology data. A.X. generated R shiny application. M.L., X.F. and S.L.B quantified stool MK concentrations. S.E.P, M.K. and P.A.M performed and analyzed metabolomics data. M.K. and J.W. performed computational analyses. M.K. and B.A.B. wrote the manuscript with input from all authors. All authors edited and commented on the manuscript.

## Methods

### Mouse husbandry

All animals were treated and housed in accordance with the Guide for Care and Use of Laboratory Animals. All experimental procedures were approved by the USC’s Institutional Animal Care and Use Committee (IACUC) and are in accordance with institutional and national guidelines. Samples were derived from animals on approved IACUC protocol number 20770 and 21212.

For the “aging cohort,” female and male C57BL/6JNia mice (4- and 20-month-old animals) were obtained from the National Institute on Aging (NIA) colony at Charles River Laboratories. For the “VCD cohort” and “fecal microbiota transplantation (FMT) cohort”, female and male C57BL/6NTac mice (3.5-month-old animals) were ordered from Taconic Biosciences. All animals were acclimated at the specific-pathogen-free animal facility at USC for 2 weeks before any processing. All animals were fed PicoLab Rodent Diet 20 (LabDiet, 5053). The facility is on a 12-h light/dark cycle and animal housing rooms are maintained at 72 °F and 30 % humidity. All animals were euthanized between 8–11 am, by CO_2_ asphyxiation followed by cervical dislocation.

For the “VCD cohort,” animals were subjected to daily intraperitoneal injections of either vehicle (safflower oil) or VCD (at a dosage of 160 mg/kg/day) for 15 consecutive days. All injections were performed between 8:00 AM and 10:00 AM to minimize variability due to circadian effects. Lethality was observed in male VCD-injected mice (4 out of 10 males in the experimental group within 16 days of the start of injections). Intriguingly all lethality cases occurred in males that had to be single-housed prior to injections due to fighting and wounds. Effects may have been aggravated due to lack of social interactions and/or temperature regulation.

### Ovarian follicle counts

Ovaries were fixed in Bouin’s solution (Sigma, HT10132) for 24 hrs at room temperature and transferred to 70 % ethanol for storage. Paraffin embedding, tissue sectioning and staining with hematoxylin and eosin (H&E) were performed by the USC Norris Comprehensive Cancer Center Translational Pathology Core Facility. H&E staining slides were imaged on the Keyence BZ-X All-in-One Fluorescence Microscope platform. Three sections per ovary were used to count the number of primordial, primary, secondary, and antral follicles and corpus luteum by three blinded observers. Median counts across the 3 observers were used for data analysis. Statistical significance between groups was assessed using the non-parametric Wilcoxon rank-sum test.

### Quantification of serum AMH, FSH and Inhibin A concentrations

After euthanasia, blood was collected directly from the heart. Blood was allowed to clot at room temperature for 1 hour, and serum was separated using MiniCollect® Serum Tube (Greiner, 450472), then stored at -80 °C until further processing. Quantification of serum AMH (Rat and Mouse Anti-Müllerian hormone (AMH) ELISA kit, Ansh Labs, AL-113), FSH (Millipore Pituitary Panel Multiplex kit, RPT86K and Ultra-Sensitive Mouse & Rat FSH, UVA Ligand Core, in-house^109^), and Inhibin A (Inhibin A ELISA kit, Ansh Labs, AL-161) levels were performed by the University of Virginia Center for Research in Reproduction Ligand Assay and Analysis Core. Standard normalized values were provided by the core. Statistical significance between groups was assessed using the non-parametric Wilcoxon rank-sum test.

For FSH quantification, two different kits were used during the course of the study, necessitating correction of the dataset generated by the ultra-sensitive kit. To ensure consistency, we generated data using both kits with the same serum samples and devised a polynomial model to fit the relationship between the two datasets (R² = 0.90). The trained polynomial model was then applied to correct the data generated by the Ultra-Sensitive Mouse & Rat FSH assay. This correction process was specifically applied to the data for the FMT cohort. The raw data used to train the model has been uploaded to Github (https://github.com/BenayounLaboratory/Ovarian_Aging_Microbiome).

### Ovarian health index calculation

The ovarian health index was calculated by integrating two key components: (1) ovarian hormone levels (AMH, FSH, and Inhibin A) and (2) follicle counts (combined counts of primordial, primary, secondary, antral follicles, and corpus luteum). A 3-tier scoring system was used for each parameter. Values beyond the young medians were assigned a score of 3, values between the young and estropausal medians were assigned a score of 2, and values beyond the estropausal medians were assigned a score of 1. The average hormone score (calculated from the AMH, FSH, and Inhibin A scores) was then combined with the follicle score in a 1:1 ratio to generate the overall ovarian health index. This combined score was scaled to a 0-100 range for ease of interpretation.

Statistical significance between groups was assessed using the Wilcoxon rank-sum test. Scripts used to calculate the ovarian health index are available on the Benayoun lab GitHub at https://github.com/BenayounLaboratory/Ovarian_Aging_Microbiome.

### DNA extraction for 16S rRNA amplicon and shotgun metagenomics analyses

Fecal matter for 16S rRNA V3-V4 amplicon and shotgun metagenomics analyses were collected aseptically post-euthanasia. Samples were immediately snap frozen in sterile tubes and stored at -80 °C. DNA extraction was performed using QIAamp Fast DNA Stool Mini Kit (QIAGEN, 51604), following the manufacturer’s protocol.

V3-V4 region targeted amplification and sequencing were performed by Novogene Inc. Amplification primers used were CCTAYGGGRBGCASCAG (FWD_341F) and GGACTACNVGGGTWTCTAAT (REV_806R). Amplicon libraries were sequenced on the Illumina NovaSeq 6000 machine. Shotgun metagenomics library preparation and sequencing were performed by Novogene Inc. on the Illumina NovaSeq X machine. De-multiplexed FASTQ files were provided by Novogene Inc. for data analysis.

### 16S rRNA V3-V4 amplicon sequencing data analysis

#### Data pre-processing, quality control and denoising

Data pre-processing, quality control and denoising were performed within the QIIME2^110^ (v. 2023.7) platform. Each cohort was analyzed independently for memory and computing efficiency. FASTQ files were imported to QIIME2 and denoised using DADA2^111^. Forward and reverse reads were truncated at 226 bp and 224 bp, respectively, to obtain reads with quality scores above 25 and to ensure coverage of the V3-V4 region (–p-trunc-len-f 226 –p-trunc-len-r 224).

#### Batch correction and principal component analysis of amplicon sequence variants

Previous studies have shown that effectiveness of batch correction tools are dataset-dependent^41^. For batch correction, a previously described benchmarking pipeline^41^ was used to select the most effective batch correction tool for our dataset. removeBatchEffect function from limma^112^ (v. 3.50.1), Combat from sva^113^ (v. 3.42.0) and FAbatch from bapred^114^ (v. 1.1) were benchmarked. Denoised feature tables were imported to R using qiime2R^115^ package (v. 0.99.6). Then, features with low counts (less than 0.01% of all the counts) were removed and filtered counts were centered log ratio (CLR)-transformed. Based on principal component analysis results (method described below) and variance calculations, FAbatch-corrected values were chosen for downstream analyses.

#### Principal component analysis of amplicon sequence variants

For principal component analysis (PCA), pca function from mixOmics^116^ (v. 6.18.1) was used in R. For female and male aging cohorts, FAbatch-corrected values were used for PCA. For VCD and FMT cohorts, CLR-transformed values were used for PCA.

#### Beta diversity and alpha diversity analyses

For beta and alpha diversity analyses, phylogenetic trees were generated using the ‘qiime phylogeny align-to-tree-mafft-fasttree’ command. Rooted phylogenetic trees were used to run the ‘qiime diversity core-metrics-phylogenetic’ command for beta and alpha diversity analyses. Diversity analysis results were imported to R using qiime2R^115^ package (v. 0.99.6) for further plot generation and significance tests. Bray-Curtis dissimilarity and Jaccard index principal coordinate analysis results were used to generate beta diversity plots using ggplot2 (v. 3.4.2). To ensure comparability across different cohorts, we normalized alpha diversity metrics within each cohort by dividing the values of each sample by the median value of the corresponding metric (e.g., observed features, Shannon entropy) within that cohort (e.g., CTL vs. VCD, FMT-YF vs. FMT-EF). This normalization allowed for combined cohort analysis. The statistical significance of differences in alpha diversity between groups was determined using the Wilcoxon rank-sum test.

#### Differential abundance analysis of microbial genera

Differential abundance analysis of microbial genera was performed using the ALDEx2^117^ plugin within the QIIME2^110^ platform. Taxonomic classifier was generated using the SILVA^118^ (132 release) reference. Reference reads were extracted for CCTAYGGGRBGCASCAG (FWD_341F) and GGACTACNVGGGTWTCTAAT (REV_806R) primer set and a classifier was trained using the ‘qiime feature-classifier extract-reads’ and ‘qiime feature-classifier fit-classifier-naive-bayes’ commands, respectively. Differential abundance analysis results and taxonomic classifier were imported to R using qiime2R^115^ for comparative analysis and plot generation. For female and male aging cohorts, differential abundance analysis results for each cohort for each sex was combined based on the directionality of the differential abundance measures, identifying features that consistently exhibited either an increase or decrease across all cohorts for each sex. Using Fisher’s method and Benjamini-Hochberg (BH) correction method, p-values for each feature’s differential abundance were combined and corrected, respectively. Additionally, the mean of effect size across cohorts for each feature was used for analysis. For combined female and aging cohorts, as well as VCD and FMT cohorts, features with an adjusted p-value < 0.05 and an absolute effect size > 1 were identified as significantly different in abundance.

#### Functional abundance prediction analysis

Functional abundance prediction analysis was performed via the PICRUSt2^48^ plugin within the QIIME2^110^ platform using default parameters. PICRUSt2 results were imported to R using qiime2R. For differential abundance and statistical significance analyses, ALDEx2^117^ was used (500 Monte Carlo samples, Wilcoxon test for statistical testing, effect size calculation enabled, and the interquartile log ratio (iqlr) for denominator calculation). To combine independent cohorts for female and male aging cohorts, Fisher’s method and Benjamini-Hochberg multiple testing correction were used to combine and correct the p-values. Also, the mean values of effect sizes were calculated for each feature. Adjusted p-value less than 0.05 and absolute effect size greater than 1 were used as criteria to filter functional pathways that were significantly different in abundance.

### Fecal microbiota transplantation

For fecal microbiota transplantation (FMT), animals underwent a 10-day course of antibiotics/antimycotic (Ab/Am) treatment before transplantation. The regimen included ampicillin in drinking water at 1 g/L (Sandoz, 00781), and oral gavage of amphotericin-B (Sigma, A9528) at 1 mg/kg/day, metronidazole (MFR VIONA PHARMACEUTICALS, 096988) at 100 mg/kg/day, vancomycin (Alvogen, Vet-Rx-MW 090160) at 50 mg/kg/day, and neomycin (MFR VETONE, 510570) at 100 mg/kg/day. Starting on the third day after the end of the Ab/Am treatment, FMT was performed biweekly for three weeks via oral gavage. Donor stool samples from more than 30 young or estropausal female mice were pooled and prepared by suspending in saline (20 mg/ml), followed by vortexing and centrifugation to obtain the supernatant. Each recipient animal received 100 µl of the fecal suspension. Two independent cohorts were enrolled in the study, each using independently prepared pools of FMT materials. Sample collection for subsequent processing and analysis occurred three days after the final FMT session.

### Ovarian RNA-seq sample preparation

Ovaries were harvested and immediately snap-frozen in sterile tubes, and then stored at -80 °C. For RNA extraction, 600 µl of Trizol reagent (Ambion, 15596018) was added to the ovaries in Lysing Matrix D tubes (MP, 6913500) and tissues were homogenized with a BeadBug 6 microtube homogenizer (Benchmark Scientific, D1036) at 3,500 rpm for 30 seconds for 3 cycles in the cold room. Total RNA was purified using the Direct-Zol RNA Miniprep (Zymo Research, R2052), following the manufacturer’s instructions. Integrity and quality of the purified total RNA were assessed on the 4200 TapeStation system (Agilent, G2991A) using High Sensitivity RNA ScreenTapes (Agilent, 5067–5579) to determine the RNA Integrity Number (RIN). Subsequently, mRNA-seq libraries were prepared and sequenced by Novogene Inc. to obtain PE150 sequencing reads on the NovaSeq X Plus platform.

### FMT cohort ovarian RNA-seq data analysis

#### Data quality control and pre-processing

Paired-end 150-bp reads were hard-trimmed to remove the first 6 bases and keep up to 100bp using Fastx Trimmer^119^ (v. 0.0.14) and processed using Trim Galore^120^ using default parameters (v. 0.6.6). Trimmed reads were mapped to the mm39 genome reference using STAR^121^ (v. 2.7.9a). Read counts were assigned to genes from the mm39 genome reference using subread^122^ (v. 2.0.3). After assigning read counts to genes, downstream processing was performed in R (v. 4.1.2). Low count genes (genes with less than 1 count in at least 6 out of 14 RNA-seq libraries) were removed to improve memory and speed efficiency of DESeq2^123^, as recommended by the developers. We used surrogate variable analysis to estimate and correct for unwanted experimental noise. R package sva^113^ (v. 3.42.0) was used to estimate surrogate variables and the removeBatchEffect function from limma^112^ (v. 3.50.1) was used to regress out the effects of surrogate variables and RNA integrity differences (RNA integrity number scores) from raw read counts. To remove extreme values within the read counts, Winsorize function from DescTools^124^ (v. 0.99.50) was used to adjust the values above the 99.9th percentile to the values at the 99.9th percentile. DESeq2^123^ package (v. 1.34.0) was used for downstream analyses of the RNA-seq data. Genes with FDR < 5% were considered statistically significant. For the multidimensional scaling (MDS) analysis, we calculated the distance between samples using the inverse of Spearman’s rank correlation coefficient (1-Rho) as the distance metric. These distances were then input into the cmdscale function in R to compute the MDS coordinates.

#### Gene ontology analysis of RNA-seq dataset

We used the GSEA^55^ approach through the phenoTest package. GO term annotations (ENS.GO.BP, ENS.GO.MF, ENS.GO.CC and GTRD gene sets) were sourced from ENSEMBL (Ensembl 108) and Molecular Signature Database^125^. The t-statistic from DESeq2^123^ was used to rank genes for functional enrichment analysis. For readability, only the top ten most-significant pathways with negative NES and top ten most-significant pathways with positive NES are shown in figures if more than that passed the FDR□<□5% significance threshold for GO terms and FDR < 10% for GTRD transcription factor targets (see main text for explanation).

#### Deconvolution analysis of immune cell type proportion

To estimate any differences in the immune cell proportions between FMT-YF and FMT-EF groups of the FMT cohort, single-cell RNA-seq reference-based deconvolution analyses were performed, using CSCDRNA^126^ and Granulator^127^. We downloaded the cell type annotated Seurat object of a publicly available murine ovarian single-cell RNA-seq dataset (Open Science Framework dataset ID 924fz)^56^. In the CSCDRNA analysis, the min.p parameter was set to 0.3 following the authors’ recommendation^126^, to efficiently exclude the 30% least significant portion of marker genes for each cell type. This threshold was chosen to increase efficiency and reduce noise. For Granulator, the qprogwc method^128^ was used with default parameters to estimate the immune cell proportions within the ovarian bulk RNA-seq dataset. Cell proportions were tested for significance using a Wilcoxon test rank-sum test.

#### Pre-processing and filtering candidate transcription factor-bound genes

For potential transcription factors identified from GSEA^55^ analysis against GTRD^61^ transcription factor targets, peak information was obtained for Cistrome.org. Peak annotation was performed using HOMER^129^ (v. 4.11.1). Genes with peaks annotated in TSS-promoter of protein coding genes were considered direct target genes. Obtained gene lists were used to perform GSEA through the phenoTest (v. 1.42.0) package.

#### Pre-processing of public ovarian aging datasets

To evaluate the correlation between ovarian aging and FMT gene expression profiles of the ovaries, we pre-processed and analyzed two publicly available ovarian aging datasets (2-3-month-old vs. 12-month-old female mice; BioProject dataset PRJNA100222 and GSA dataset CRA003645)^68,69^. For each dataset, reads were pre-processed and aligned to the mm39 genome reference using the same methods as for the FMT cohort ovarian RNA-seq dataset. Low count genes (genes with less than 1 count in at least 4 out of 7 RNA-seq libraries) were removed. sva^113^ (v. 3.42.0) was used to estimate surrogate variables and the removeBatchEffect function from limma^112^ (v. 3.50.1) was used to regress out the effects of surrogate variables from raw read counts. DESeq2^123^ package (v. 1.34.0) was used for downstream analyses of the RNA-seq data. p-values for common genes between the two datasets were combined and corrected using Fisher’s method and BH correction method, respectively. FDR < 1e-5 was used to catalog gene sets that are robustly up- or down-regulated with aging. Obtained gene lists were used to perform GSEA through the phenoTest package.

### Fertility assay

Fertility assay was performed as previously described^71^. FMT female mice were paired with 4-month-old males at a 1:1 ratio 3 days after the last day of FMT administration until the birth of the first litter. The number of pups in the first litter and the latency, defined as the duration between the pairing date and the birth date of the first litter, were recorded and analyzed. Statistical difference in latency in the 1^st^ litter after pairing was evaluated using a log-rank test. For pairs that did not produce pups within the 60-day timeframe, the latency was recorded as 60 days.

### Whole genome shotgun metagenomics analysis

#### Data quality control and pre-processing

Paired-end 150-bp reads were pre-processed to remove sequencing adapters and low-quality reads using Trimmomatic (v. 0.39)^130^. Trimmed reads were then mapped to the mouse reference genome (mm10) using Bowtie2 (v. 2.4.4)^131^ to remove host DNA contamination. Unmapped reads from paired-end files were extracted using the --un-conc-gz option in Bowtie2 and used for subsequent analyses. Quality-controlled and host-depleted reads were then processed for taxonomic classification and functional profiling.

#### Taxonomic classification, abundance estimation and functional profiling

Kraken2 (v. 2.1.2)^132^ was used for taxonomic classification of unmapped reads. Reads were classified against a pre-built standard Kraken2 database containing bacterial, archaeal, and viral sequences. Classification outputs were summarized using Bracken (v. 2.7)^133^ to refine species- and genus-level abundance estimates. Bracken was configured with a read length of 150 bp and default settings for abundance profiling. Functional profiling was conducted using HUMAnN3 (v. 3.0)^74^. Reads were processed in paired-end mode. Pathway and gene family tables were further merged and normalized using human_join_tables and human_renorm_table functions.

#### Alpha and beta diversity and differential abundance analysis

Raw taxonomic abundance tables from Bracken (v. 2.7)^133^ were loaded in R and filtered to remove low-abundance and low-prevalence features. Features with total counts <20, relative abundance <0.01%, or prevalence in fewer than 10% of samples were excluded. Samples with high host DNA alignment rates (i.e., MHK271) were also removed. Centered log-ratio (CLR) transformation of filtered relative abundances was performed. Shannon entropy was calculated using the mia^134^ (v. 1.12.0) R package. CLR-transformed relative abundance data were analyzed using mixOmics^116^ (v. 6.28.0) and Bray-Curtis and Jaccard distance matrices were calculated using vegan (v. 2.6-8). Differential abundance of species was assessed using ALDEx2^117^ (v. 1.36.0). Features with adjusted p-values of <0.05 were deemed significant.

#### Functional pathway analysis

Relative abundance data for microbial pathways from HUMAnN3^74^ (v. 3.9) was loaded in R and low-abundance pathways with total abundance below 0.001 across all samples were removed. Differential abundance of microbial pathways between groups was analyzed using ANCOMBC2^135^ (v. 2.0.0). Significant pathways with Q-values below 0.1 were identified, and log2-fold change values were used to assess the direction and magnitude of differential abundance.

### Stool menaquinone concentration quantification

Frozen stool samples were shipped to the Vitamin K Laboratory at the Tufts University Human Nutrition Research Center on Aging (HNRCA), where stool menaquinones concentrations were measured using the LC/MS, as detailed elsewhere^136^.

### Serum untargeted metabolomics

#### Sample processing

Serum samples were stored at -80°C until they were thawed on ice for analysis. Metabolites were extracted by adding 80 µL of ice-cold methanol to 20 µL of serum, followed by incubation on ice with vortexing every 5 minutes. The samples were then centrifuged at 21,300 × g for 10 minutes at 4°C.

#### Liquid Chromatography-Tandem Mass Spectrometry (LC-MS/MS)

Untargeted metabolomics was performed at the USC Alfred E. Mann Multi-Omics Mass Spectrometry Core. A volume of 10 µL from each sample was injected onto a SeQuant ZIC-pHILIC Column (5 µm, 2.1 mm × 150 mm; Millapore Sigma) using a Vanquish Flex UHPLC System (Thermo Fisher Scientific). Chromatography was conducted at a flow rate of 100 µL/min with an optimized gradient of solvent A (water/20 mM ammonium carbonate, pH 9.8) and solvent B (acetonitrile) as follows: 0 min/97% B, 1 min/97% B, 28 min/75% B, 39 min/50% B, 44 min/20% B, 45 min/20% B, 45 min/97% B, 60 min/97% B. The chromatography system was coupled to a heated electrospray ionization (HESI) source and a hybrid quadrupole-Orbitrap mass spectrometer (Q Exactive, Thermo Fisher Scientific). Mass spectra were acquired in Full MS/data-dependent MS2 mode, alternating between positive and negative ion polarities.

Source parameters were set as follows: a source voltage of 3.5 kV, source temperature of 275°C, sheath gas of 45, auxiliary gas of 10, sweep gas of 2, and capillary temperature of 320°C. Full scans were performed across an m/z range of 70 - 1050 with the following settings: an automated gain control (AGC) target of 5 × 10□, a maximum injection time of 100 ms, and a resolution of 70,000 at m/z 200. Up to five consecutive MS/MS scans were acquired per full scan with the following parameters: quadrupole isolation of charge states 1–7, isolation window of 0.6 m/z, normalized collision energies of 20, 30, and 40, dynamic exclusion of 30 s, AGC target of 1 × 10□, maximum injection time of 100 ms, and resolution of 17,500 at m/z 200.

#### Data analysis

Raw data were processed using Compound Discoverer (Version 3.3 SP3, Thermo Fisher Scientific) for metabolite identification, retention time alignment, and unknown compound detection across all samples. Metabolites were identified by matching exact masses (delta ppm of >10 or <-10) and fragmentation spectra to the mzCloud database (ddMS2; ‘mzCloud Best Match’ > 50). For quality control, only ions with a peak rating > 4 in at least six samples within a group were retained. VSN normalized filtered metabolic matrix via normalizeVSN function of limma^112^ (v. 3.50.1) was used for PCA analysis. The filtered metabolic matrix was then scaled, and Linear Discriminant Analysis (LDA) was performed via the MASS (v. 7.3-60.2) R package. Metabolites with a consistent direction of log_₂_ fold change (log_₂_FC) between two batches were subsequently retained for abundance analysis.

### Causal mediation analysis

Causal mediation analysis to identify specific features and amplicon sequence variants (ASVs) from WGS and 16S rRNA amplicon sequencing datasets from the FMT cohort acting as mediators between the treatment (FMT-YF vs. FMT-EF) and outcome (changes in the ovarian transcriptome profiles) was performed using mediation package^137^ (v. 4.5.0) in R. Feature table, taxonomic reference and differential abundance analysis results generated from QIIME2^110^ were imported to R using qiime2R^115^. ASVs identified as significant in differential abundance analyses were tested. CLR-transformed ASV counts were used for the mediation analyses. For the outcome variable, the first dimension from the multidimensional scaling (MDS) analysis of the FMT ovarian mRNAseq dataset was used. Linear models were constructed for both the mediator and the outcome, incorporating the treatment as an independent variable in each. The mediate function from the mediation package was then employed to estimate the Average Causal Mediation Effects (ACME), Average Direct Effects (ADE), and Total Effects.

### Quantification and statistical analysis

All statistical analysis was performed using the R software, version 4.2.4. For all boxplots, the data is shown with the median, the 25th and 75th percentile of the data, and the whiskers represent 1.5 _∗_ the inter-quartile range (IQR). Individual datapoints are always overlayed for transparency and rigor. Specific statistical tests used, the number of biological replicates and how many animals they are indicated in the corresponding figure legends and associated method section.

## Data availability

The raw FASTQ files have been deposited to the Sequence Read Archive under accession PRJNA1076509. Raw microscopy pictures of ovarian hematoxylin and eosin staining are available on Figshare (Figshare DOIs: 10.6084/m9.figshare.25749588, 10.6084/m9.figshare.25749594, 10.6084/m9.figshare.25749591, 10.6084/m9.figshare.28462274). Processed serum metabolomics data can be found in Extended Data Table 4. Raw data can be found on Metabolomics Workbench, project ST003750. All R code was run using R 4.1.2. Any additional information required to reanalyze the data reported in this work paper is available from the corresponding author upon request.

## Code availability

All scripts used to analyze the datasets are available on the Benayoun lab GitHub at https://github.com/BenayounLaboratory/Ovarian_Aging_Microbiome.

## Extended Data Fig. legend

**Extended Data Fig. 1. Characterization of ovarian health state of young and estropausal female mice.** (A) Schematic diagram of ovarian health index calculation. (B) Representative images of ovarian hematoxylin-eosin stained sections. Scale bar: 50 µm. (C) Boxplots of ovarian follicle counts per section. The schematic diagrams of the follicles were created using the Generic Diagramming Platform (GDP)^138^. (D) Boxplots of serum concentrations of FSH and Inhibin A. Significance in nonparametric two-sided Wilcoxon rank-sum tests are reported for (C) and (D). YF: Young female; EF: Estropausal female.

**Extended Data Fig. 2. Characterization of fecal microbial profiles of young and estropausal female mice.** (A) Principal coordinate analysis results of Bray-Curtis dissimilarity (top panel) and Jaccard (bottom panel) indices. (B) Boxplots of base abundance of differentially abundant microbial genera. YF: Young female; EF: Estropausal female.

**Extended Data Fig. 3. Characterization of fecal microbial profiles of young and estropausal female mice.** Boxplots of base abundance of differentially abundant microbial genera. YF: Young female; EF: Estropausal female.

**Extended Data Fig. 4. Characterization of ovarian health state of VCD cohort mice.** (A) Schematic diagram of the experimental setup of VCD cohort. (B) Representative images of ovarian hematoxylin-eosin stained sections. Scale bar: 50 µm. (C-F) Boxplots of (C) ovarian follicle counts per section, (D) combined follicle counts, (E) ovarian health index, and (E) serum concentrations of AMH, FSH and Inhibin A of control and VCD-injected mice. The schematic diagrams of the follicles were created using the Generic Diagramming Platform (GDP)^138^. Significance in nonparametric two-sided Wilcoxon rank-sum tests are reported for (C-F).

**Extended Data Fig. 5. Characterization of fecal microbial profiles of VCD cohort mice.** (A) Principal component analysis result of CLR-transformed and batch-corrected ASV counts from control and VCD-injected mice. (B) Principal coordinate analysis results of Bray-Curtis dissimilarity (top panel) and Jaccard (bottom panel) indices. (C, D) Boxplots of normalized observed features and Shannon entropy indices of (B) females, and (D) males. Median from control-injected mice was used to normalize indices. (E) Differential abundance analysis results of microbial genera of control vs. VCD-injected female mice using ALDEx2. (F) Functional abundance prediction analysis of control vs. VCD-injected female mice using PICRUSt2. Significance in nonparametric two-sided Wilcoxon rank-sum tests are reported for (C,D).

**Extended Data Fig. 6. Differential abundance analysis of 16S amplicon sequencing data of VCD cohort female mice.** Boxplots of base abundance of differentially abundant microbial genera between control and VCD-injected female mice.

**Extended Data Fig. 7. Characterization of fecal microbial profiles of young and old male mice.** (A) Schematic diagram of the experimental setup of male aging cohorts. (B) Principal component analysis result of CLR-transformed and batch-corrected ASV counts from young and old male mice. Animals from three independent cohorts, n=14 per group (variation due to animal death or abnormal health issues prior to experiment). (C) Principal coordinate analysis results of Bray-Curtis dissimilarity (top panel) and Jaccard (bottom panel) indices. (D) Boxplots of observed features and Shannon entropy indices of young and old male mice. Medians from young males for each cohort were used to normalize indices per cohort. (E) Differential abundance analysis results of microbial genera of young and old male mice using ALDEx2. (F) Functional abundance prediction analysis of young vs. old male mice using PICRUSt2. Significance in nonparametric two-sided Wilcoxon rank-sum tests are reported for (D). YM: Young male; OM: Old male.

**Extended Data Fig. 8. Differential abundance analysis of 16S amplicon sequencing data of young and old male mice.** Boxplots of base abundance of differentially abundant microbial genera between young and old male mice.

**Extended Data Fig. 9. Bulk RNA-seq analysis of ovaries from young and estropausal female mice fecal microbiota transplantation recipient mice.** (A) Heat map of significant (FDR < 5%) genes differentially expressed in FMT-YF vs. FMT-EF mice. (B) Dot plots of immune cell proportions in percentages between young and estropausal fecal microbiota transplantation recipient mice (FMT-YF and FMT-EF, respectively) predicted by deconvolution analyses using CSCDRNA (left panel) and Granulator (right panel). (C) GSEA enrichment score plots Ncoa1-peak (left panel) and Usp7-peak (right panel) genes. (D) Bar plots of normalized log_2_ counts of *Ncoa1* and *Usp7* expression.

**Extended Data Fig. 10. Characterization of ovarian health state of FMT cohort mice.** (A) Representative images of ovarian hematoxylin-eosin stained sections. Scale bar: 50 µm. (B-E) Boxplots of (B) combined ovarian follicle counts, (C) ovarian follicle counts per section, (D) serum concentrations of AMH, FSH (Multiplex assay), FSH (Ultra-sensitive assay), FSH combined and INHBA, and (E) weight percentage difference between day 0 and 16 of FMT regimen. The schematic diagrams of the follicles were created using the Generic Diagramming Platform (GDP)^138^.

**Extended Data Fig. 11. Characterization of fecal microbial profiles of FMT cohort mice.** (A) Principal component analysis result of CLR-transformed and batch-corrected counts from young and estropausal female mice fecal microbiota transplantation recipient mice (FMT-YF and FMT-EF, respectively). (B) Principal coordinate analysis results of Bray-Curtis dissimilarity (left panel) and Jaccard (right panel) indices. (C) Boxplots of observed features and Shannon entropy indices of FMT-YF and FMT-EF mice. (D) Differential abundance analysis results of microbial species of FMT-YF vs. FMT-EF mice using ALDEx2. (E) Venn diagram of overlapping microbial genera between genera that increase in abundance with female aging (“UP in EF”) and increase in abundance in FMT-EF recipient mice (“UP in FMT-EF”). (F) Functional abundance prediction analysis of FMT-YF vs. FMT-EF mice using PICRUSt2. Significance in nonparametric two-sided Wilcoxon rank-sum tests are reported for (C). YF: Young female; EF: Estropausal female.

**Extended Data Fig. 12. Differential abundance analysis of 16S amplicon sequencing data of FMT cohort mice.** Boxplots of base abundance of differentially abundant microbial genera between FMT-YF and FMT-EF mice.

**Extended Data Fig. 13. Differential abundance analysis of WGS data of FMT cohort mice.** Boxplots of base abundance of differentially abundant microbial species between FMT-YF and FMT-EF mice.

**Extended Data Fig. 14. Stool vitamin K quantification of FMT cohort mice.** (A) Stool concentrations of combined vitamin K (MK6-MK13) of FMT-YF and FMT-EF. (B) Fold change in stool concentrations of vitamin K (MK6-MK13) of FMT-YF and FMT-EF.

**Extended Data Fig. 15. Serum untargeted metabolomics analysis of FMT cohort mice.** Boxplots of metabolite abundances of top 15 metabolites contributing to LDA separation between FMT-YF and FMT-EF.

**Extended Data Fig. 16. Causal mediation analysis of microbial species and ovarian transcriptome changes in fecal microbiota transplantation recipient mice.** (A) Graphical representation of causal mediation analysis. (B) Causal mediation analysis results of 16S amplicon ASVs that showed significant mediation effects on changes in the ovarian transcriptome in fecal microbiota transplantation recipient mice. Nonparametric bootstrap confidence intervals with the percentile method are reported. ACME: Average Causal Mediated Effect; ADE: Average Direct Effect.

## Extended Data Table

Extended Data Table 1. Weight measurements of FMT cohort.

Extended Data Table 2. Estrous cycle staging data of FMT cohort.

Extended Data Table 3. ALDEx2 differential abundance analysis result of metagenomics data of FMT cohort.

Extended Data Table 4. MS2-filtered metabolite abundance matrix.

