## Extended Data Figures for "Estropausal gut microbiota transplant improves measures of ovarian function in adult mice"

Extended Data Fig. 1

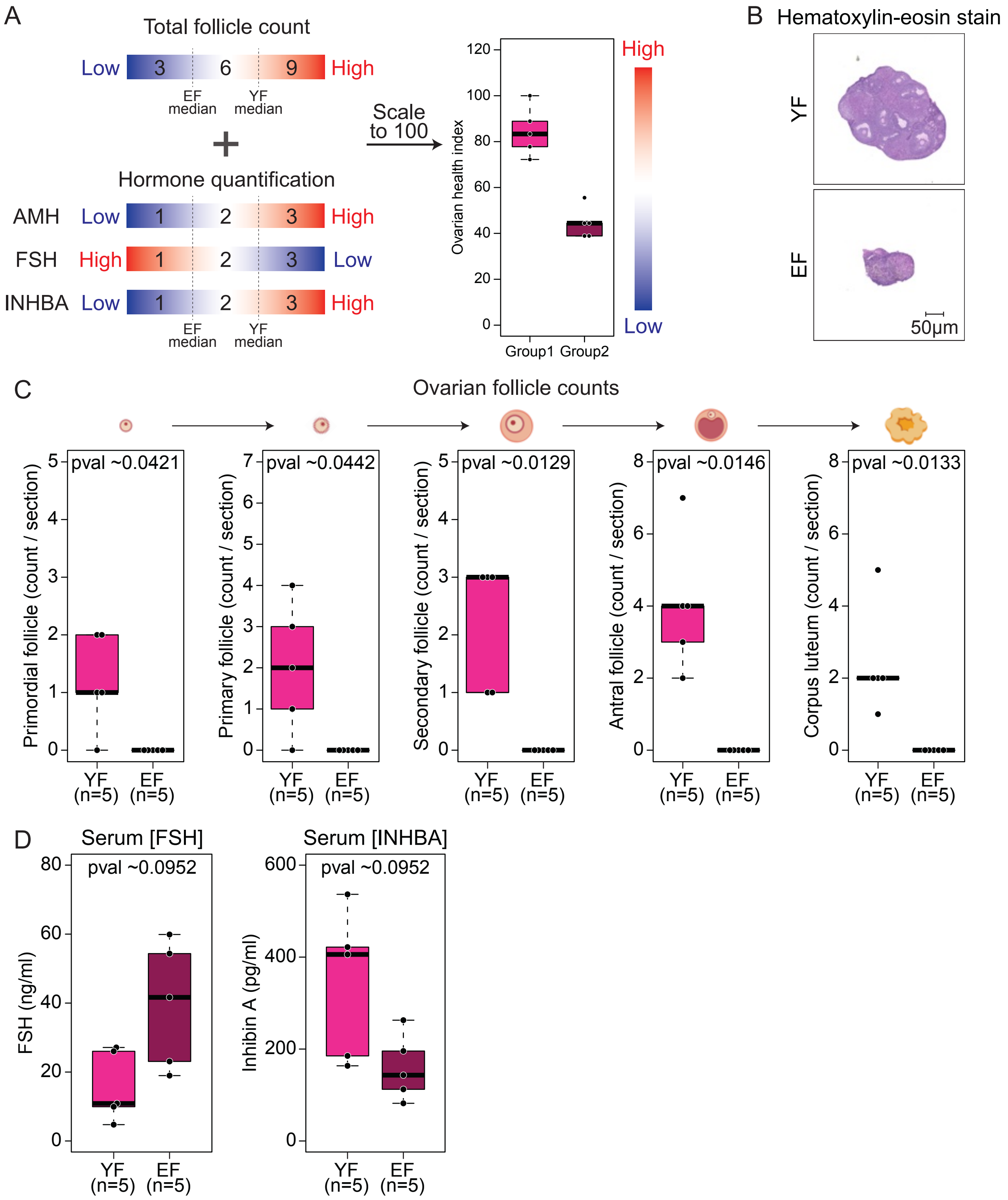

Extended Data Fig. 2

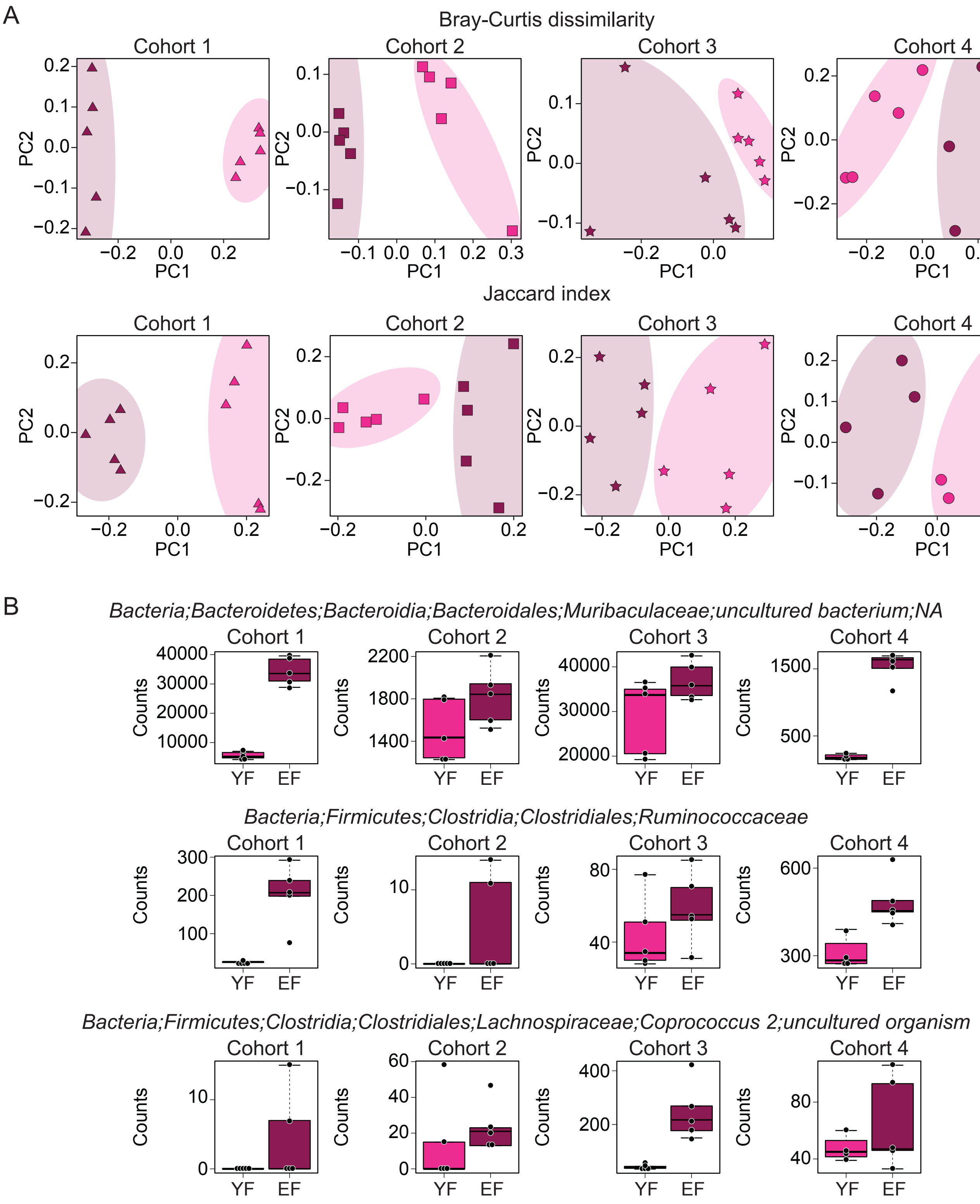

Extended Data Fig. 3

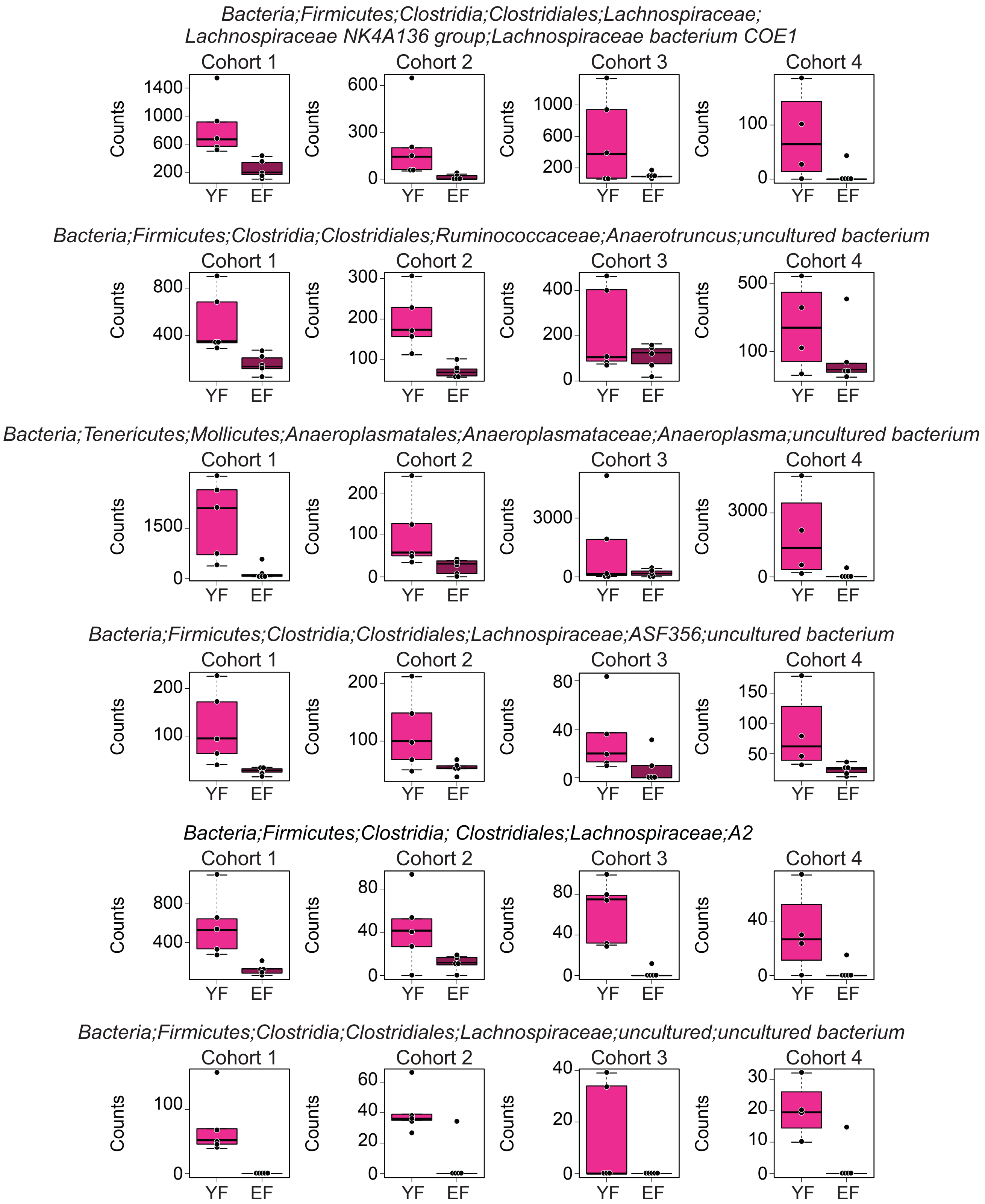

Extended Data Fig. 4

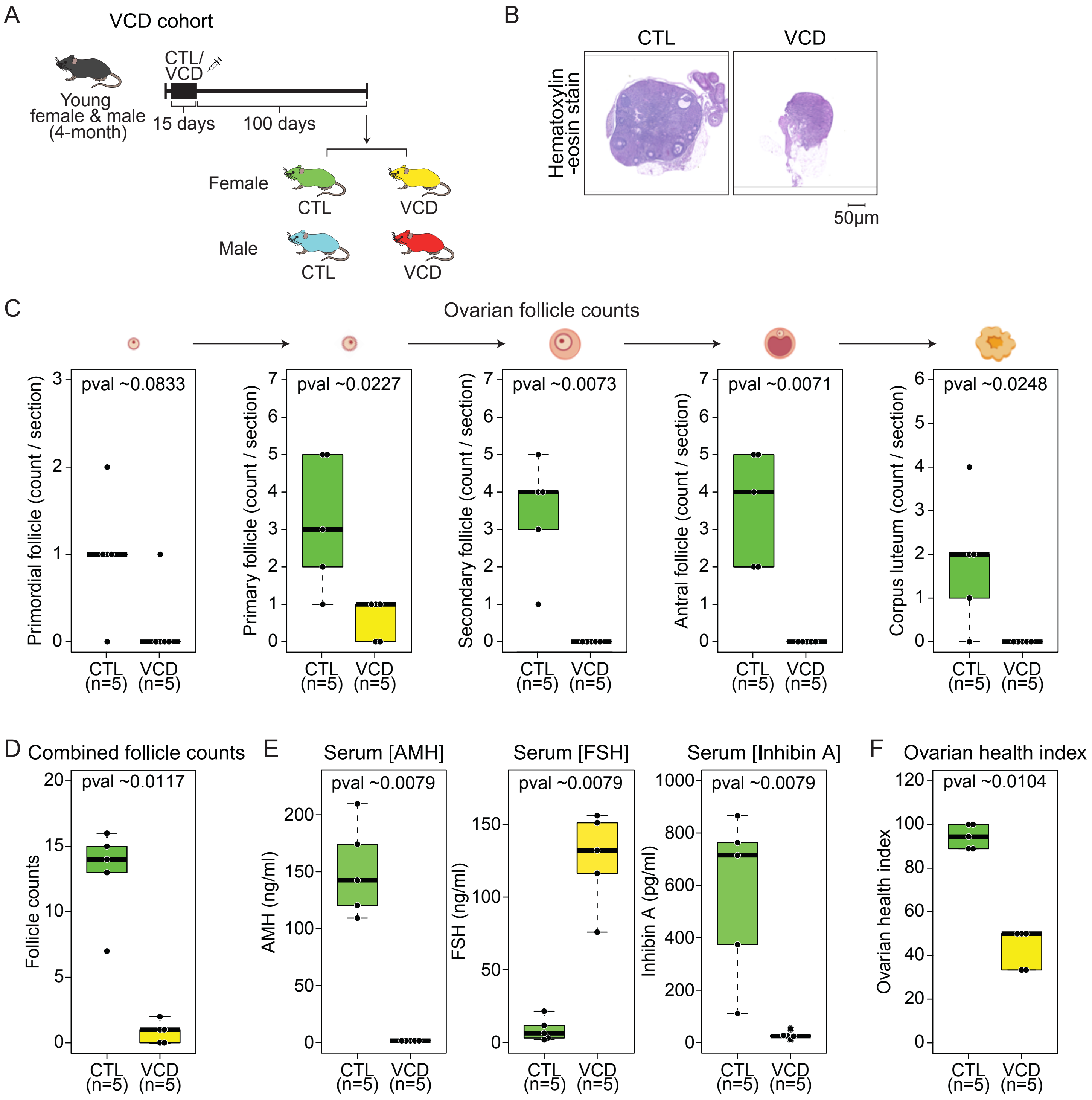

Extended Data Fig. 5

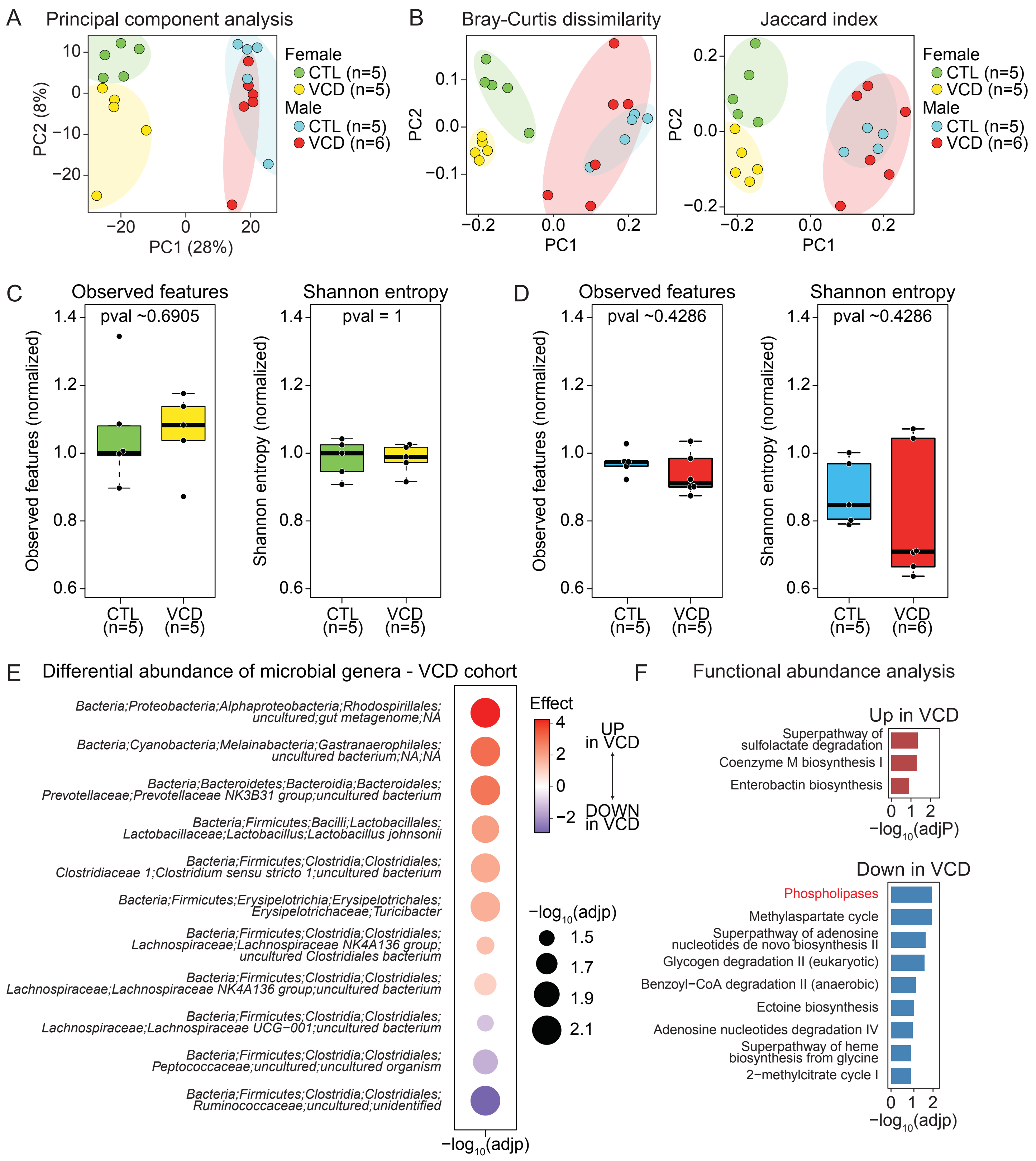

Extended Data Fig. 6

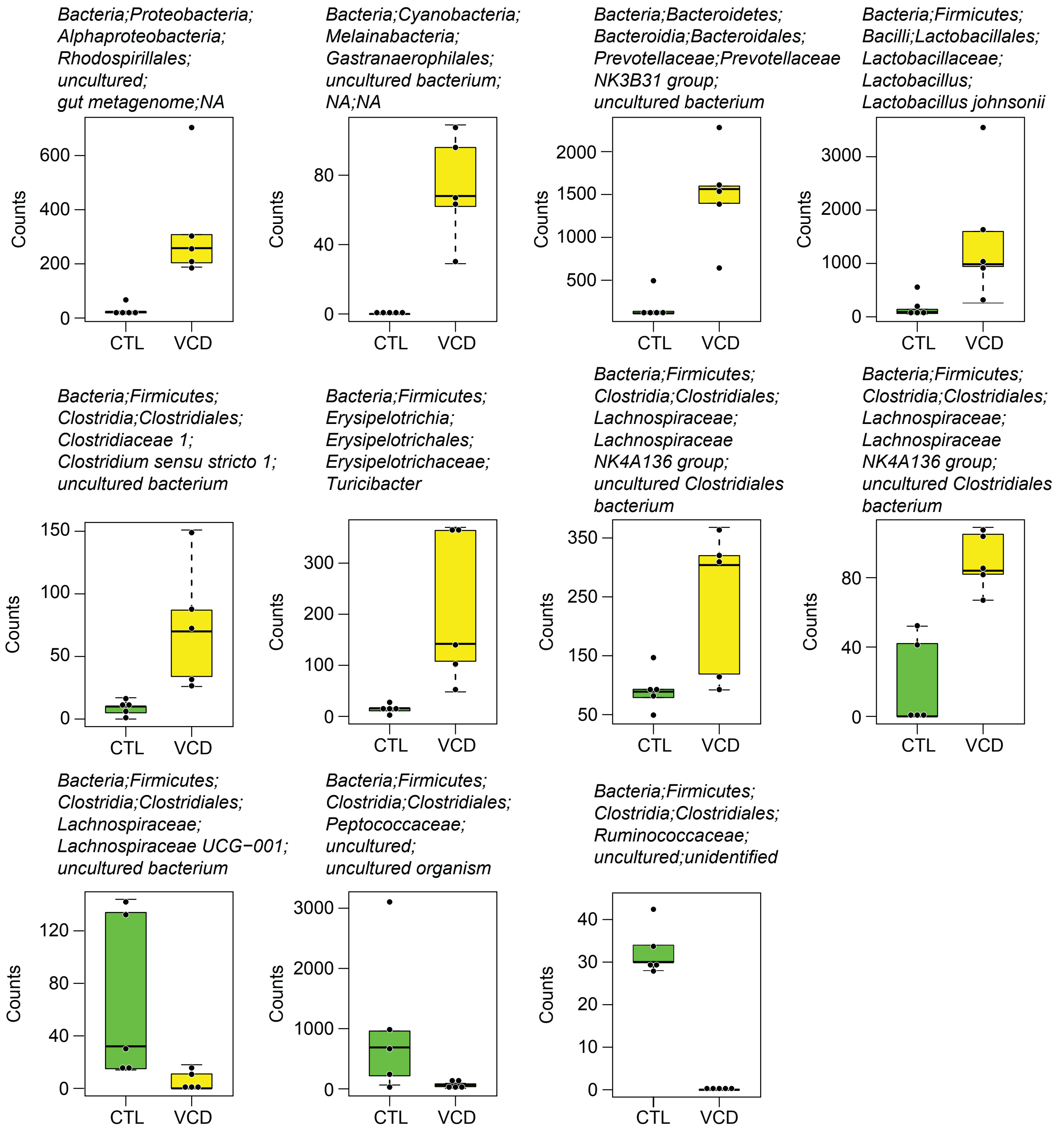

Extended Data Fig. 7

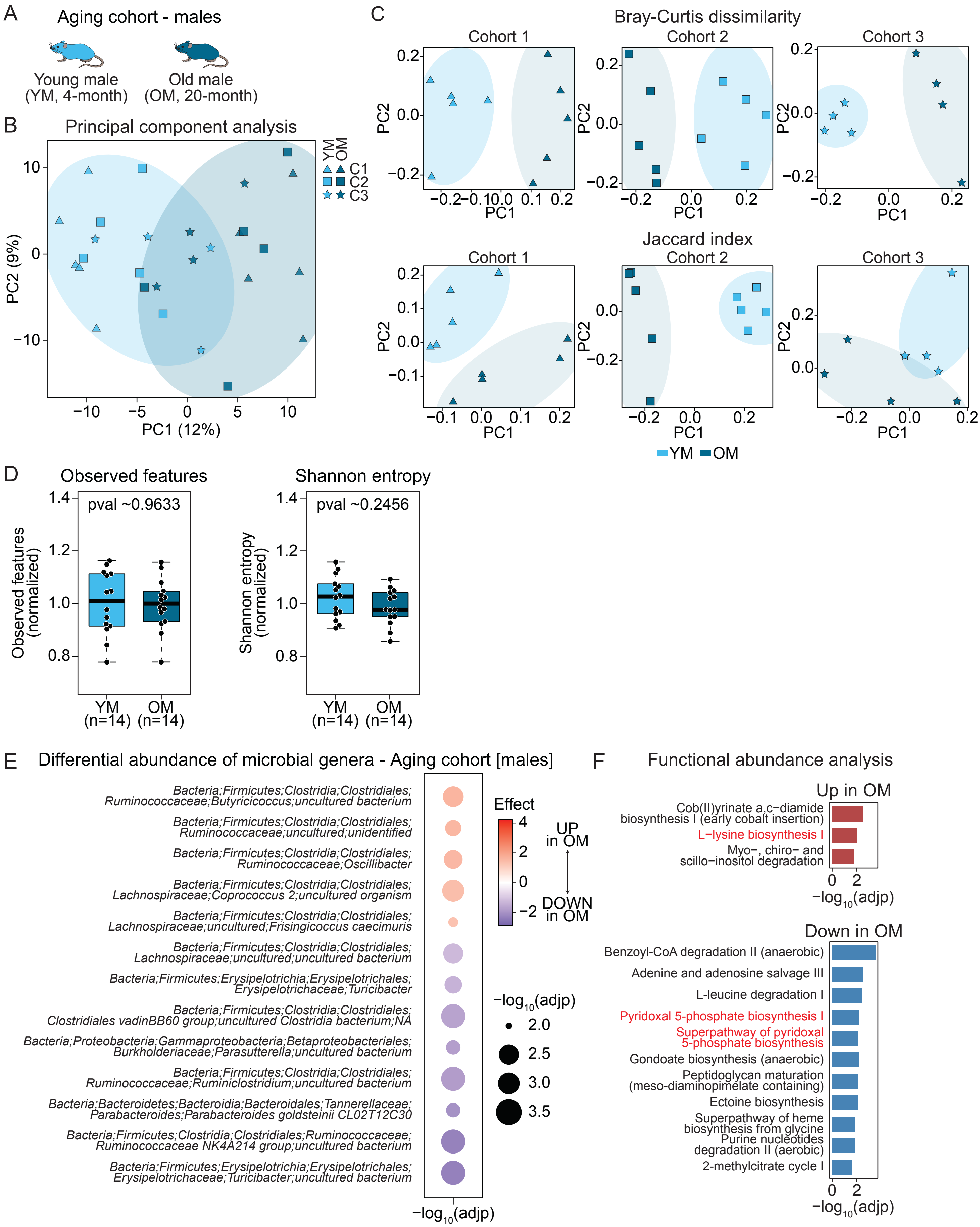

Extended Data Fig. 8

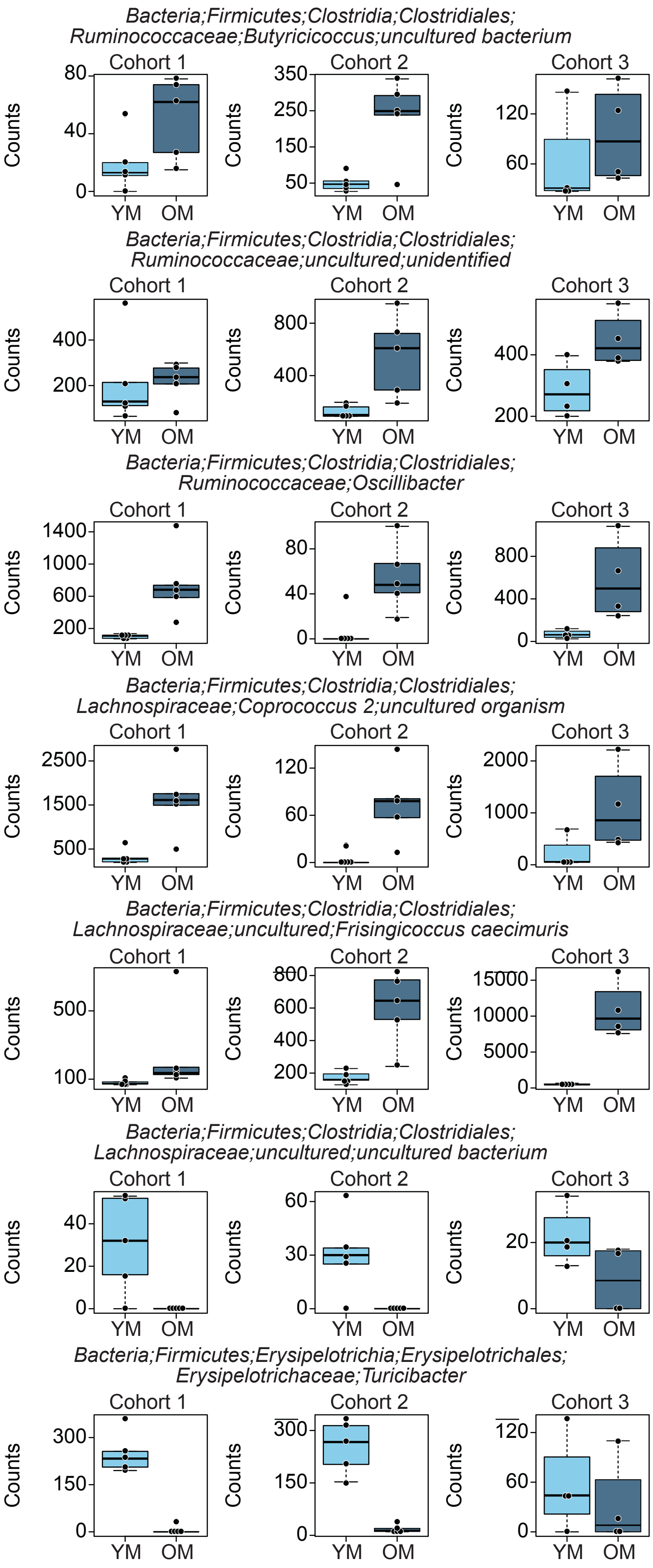

Extended Data Fig. 9

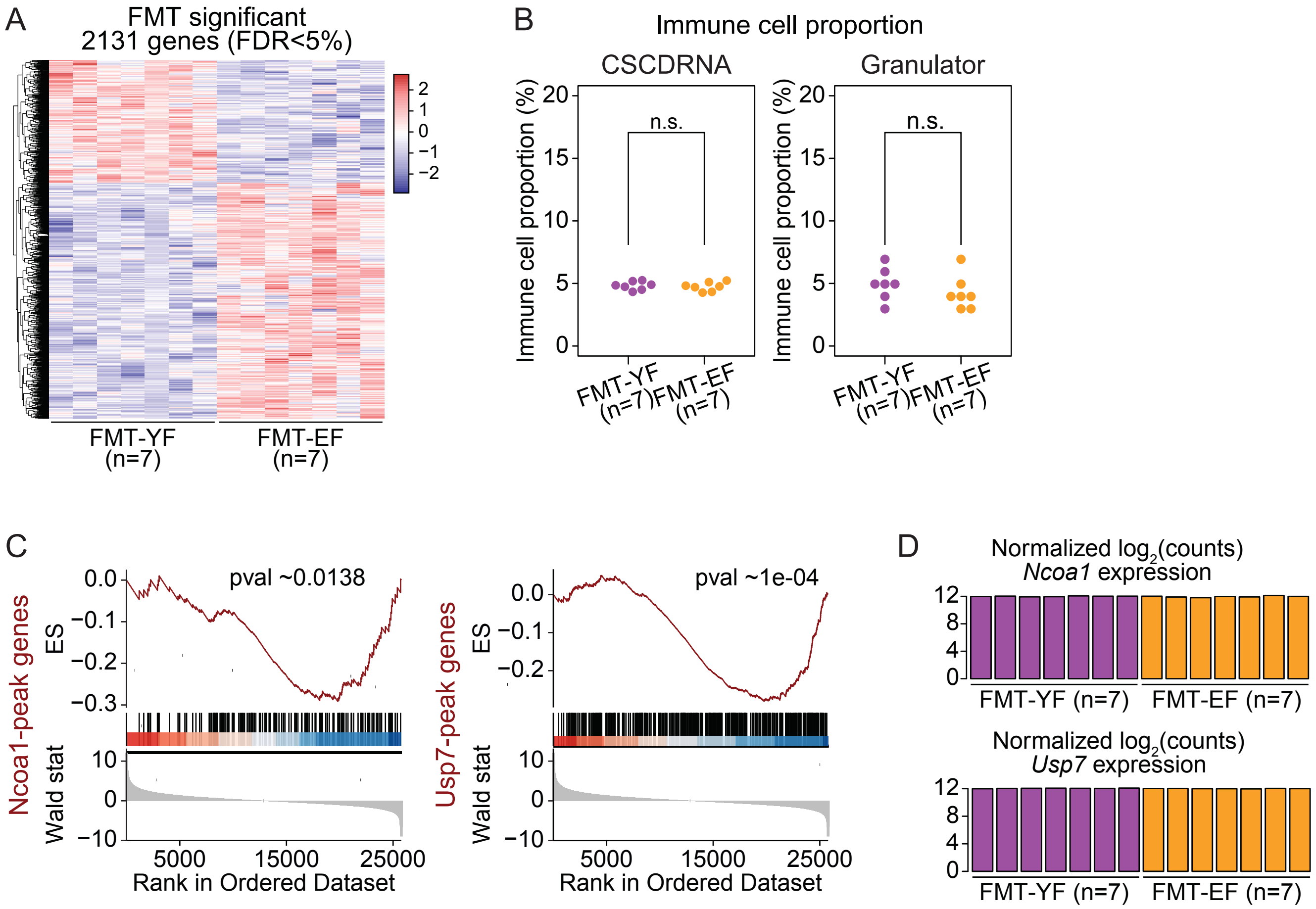

Extended Data Fig. 10

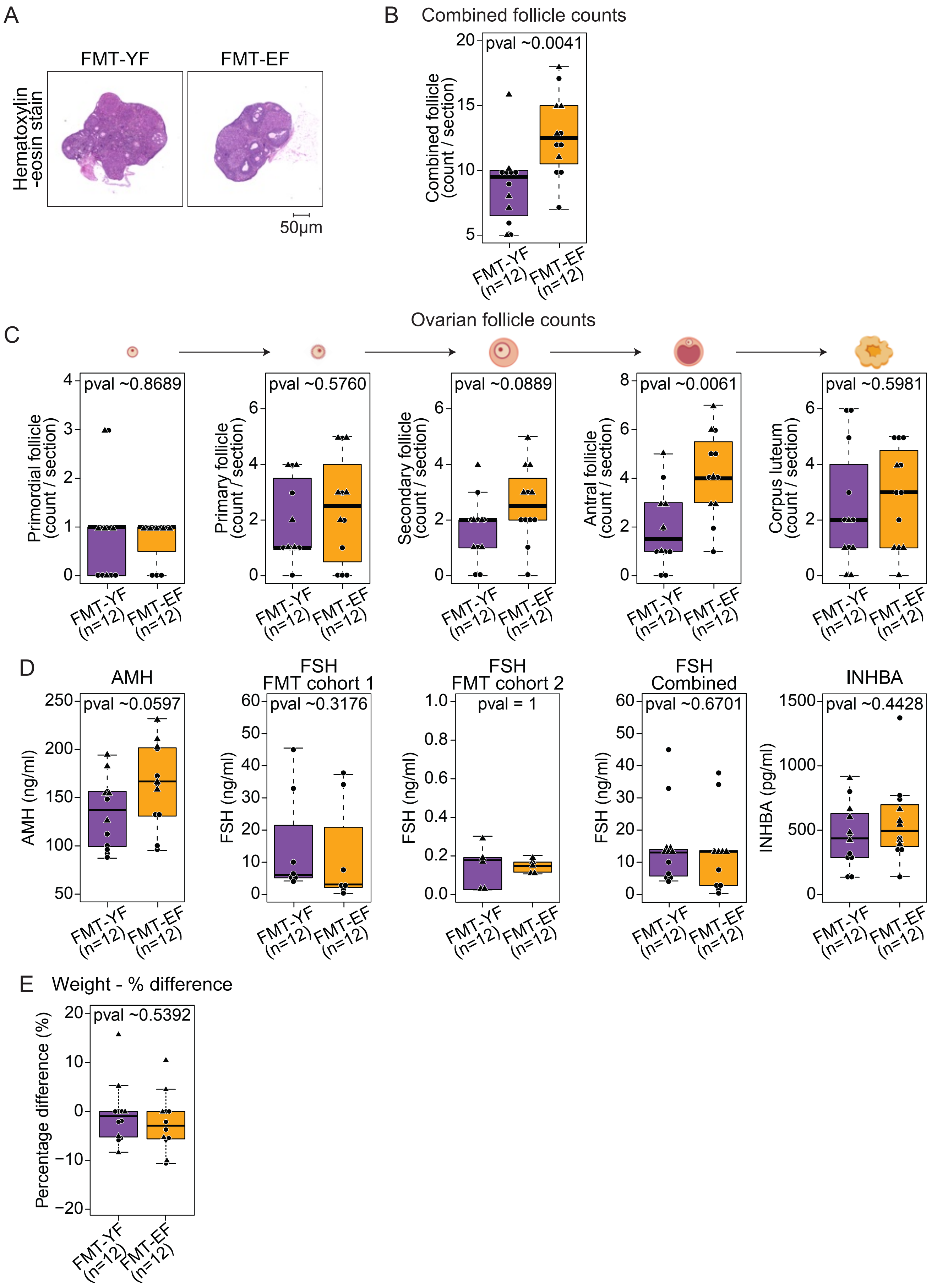

Extended Data Fig. 11

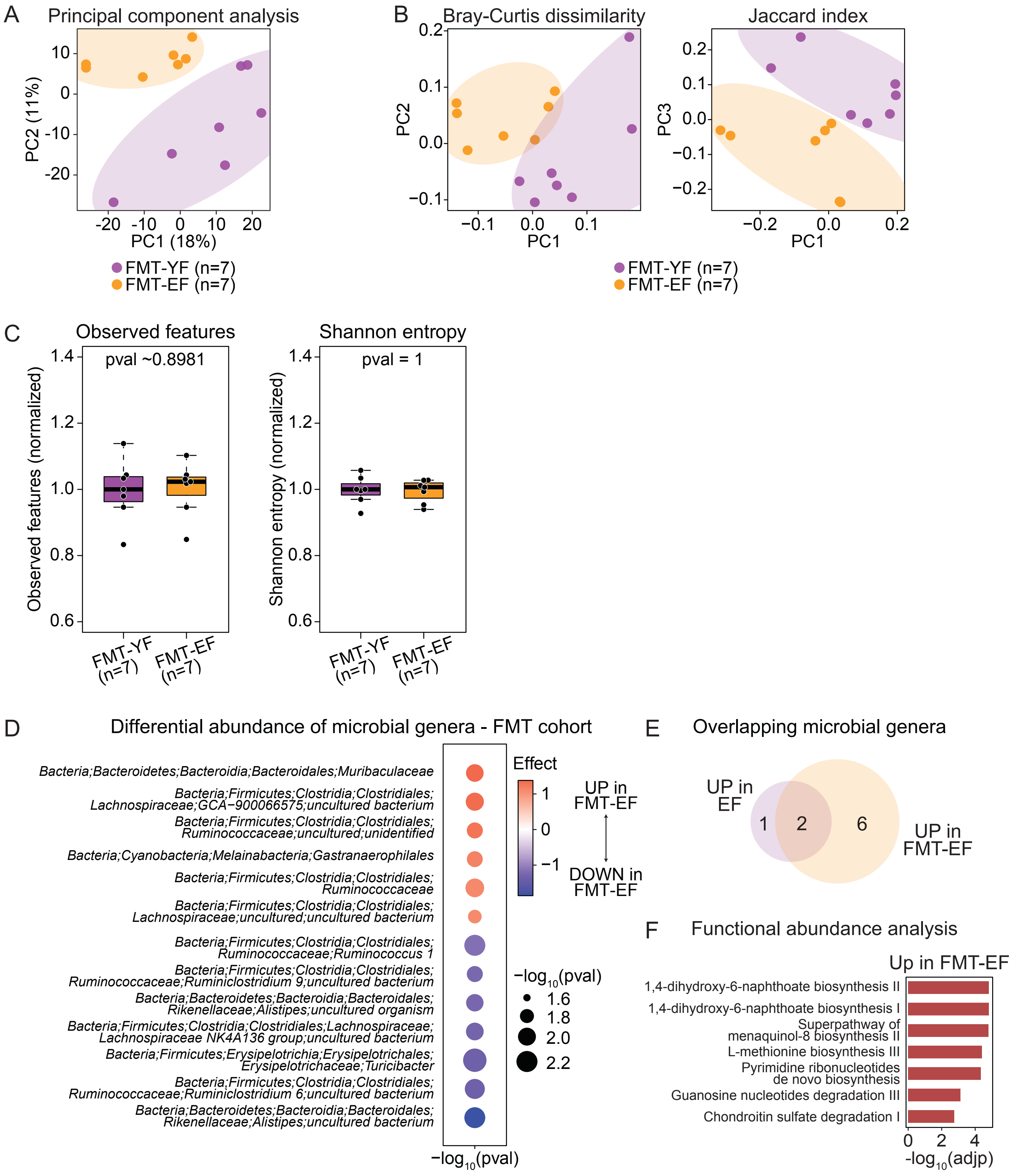

Extended Data Fig. 12

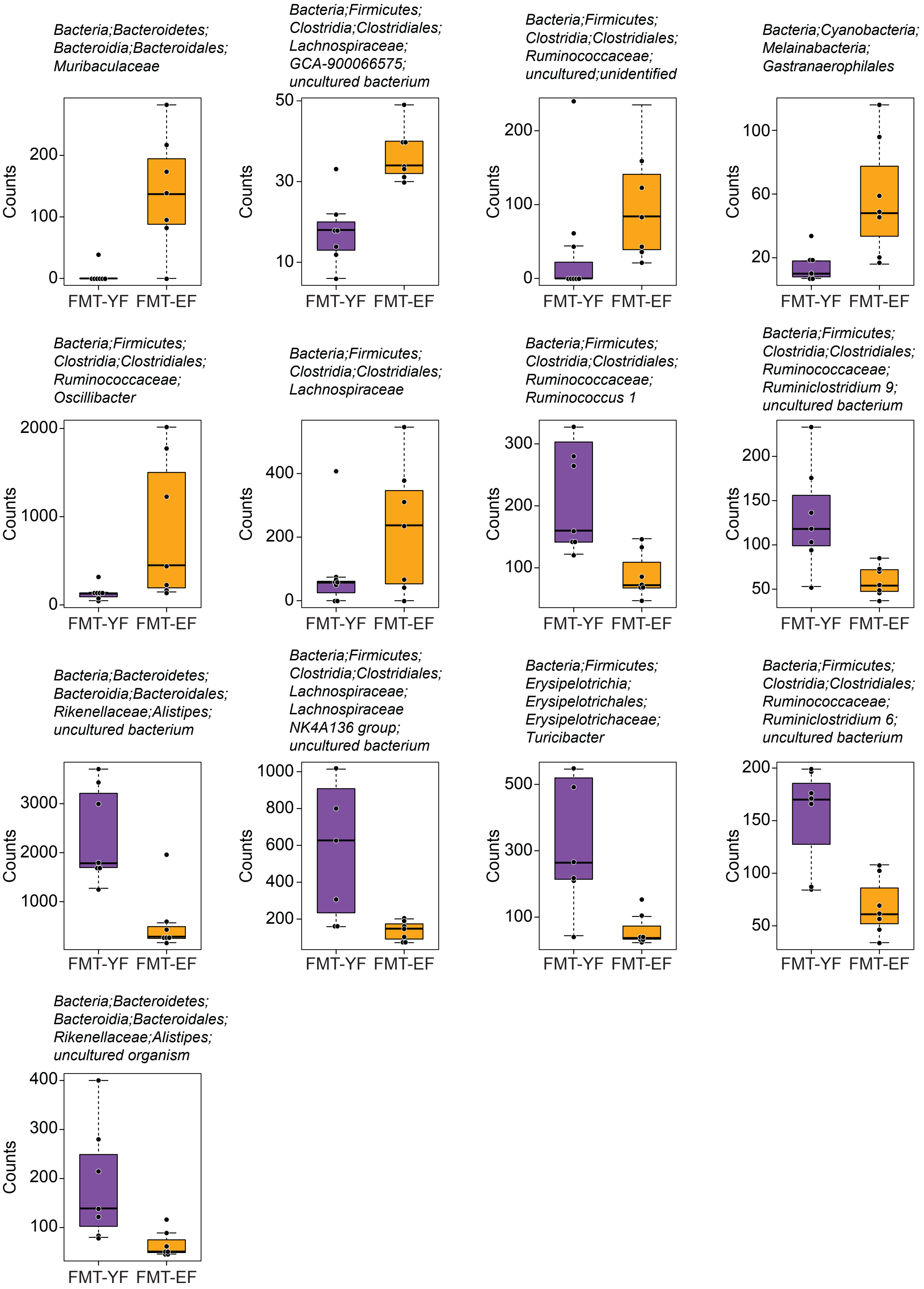

Extended Data Fig. 13

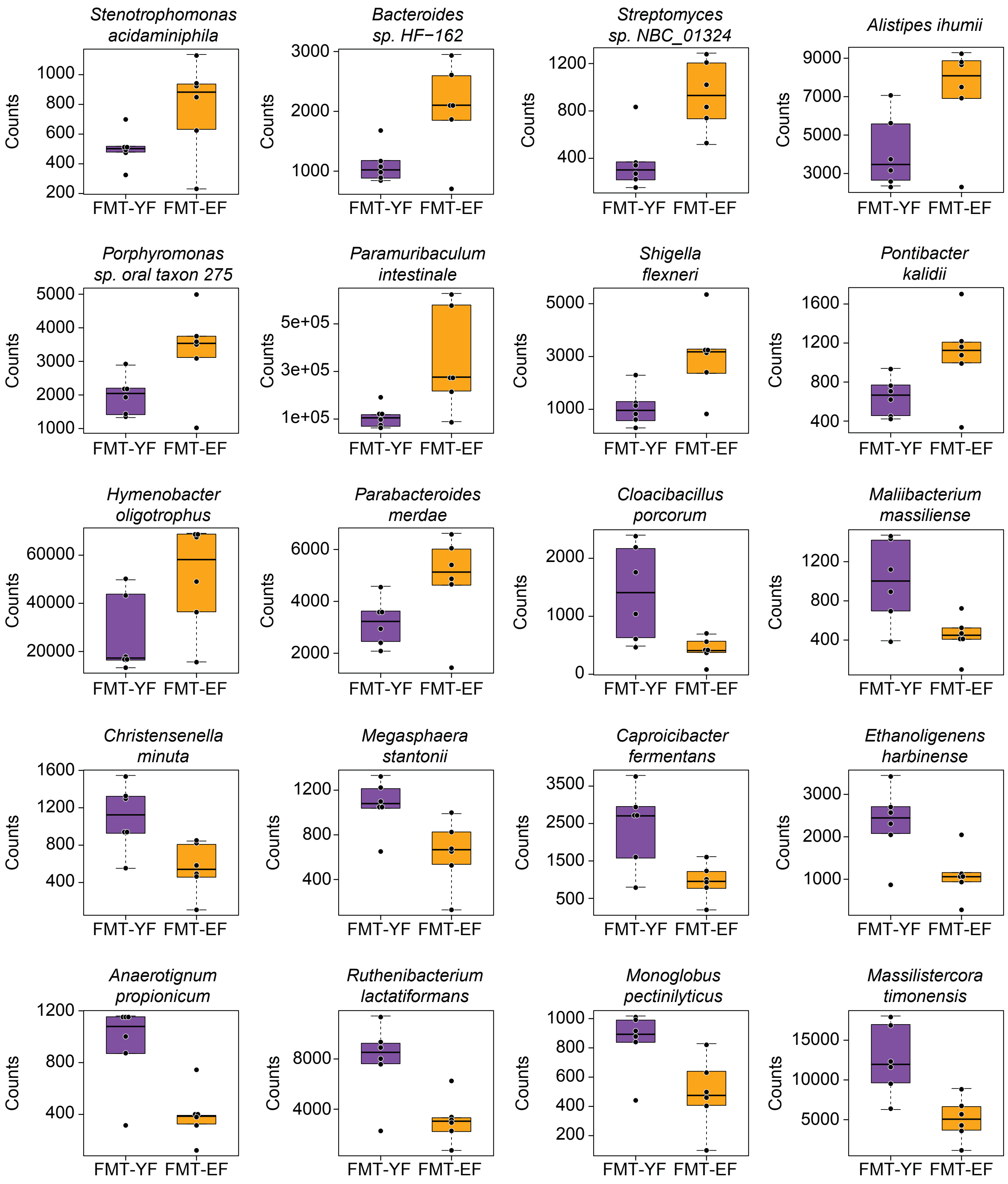

Extended Data Fig. 14

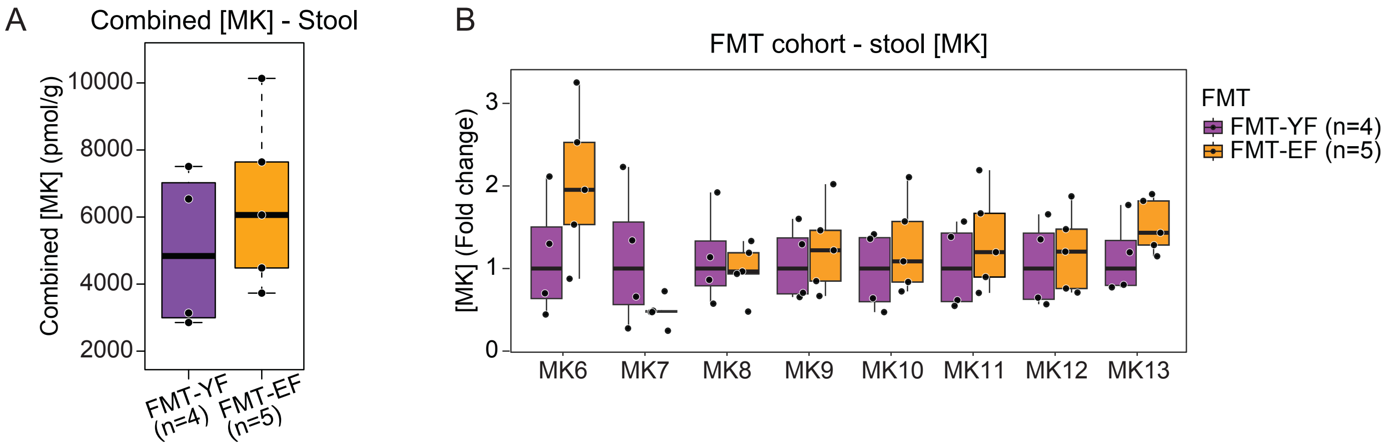

Extended Data Fig. 15

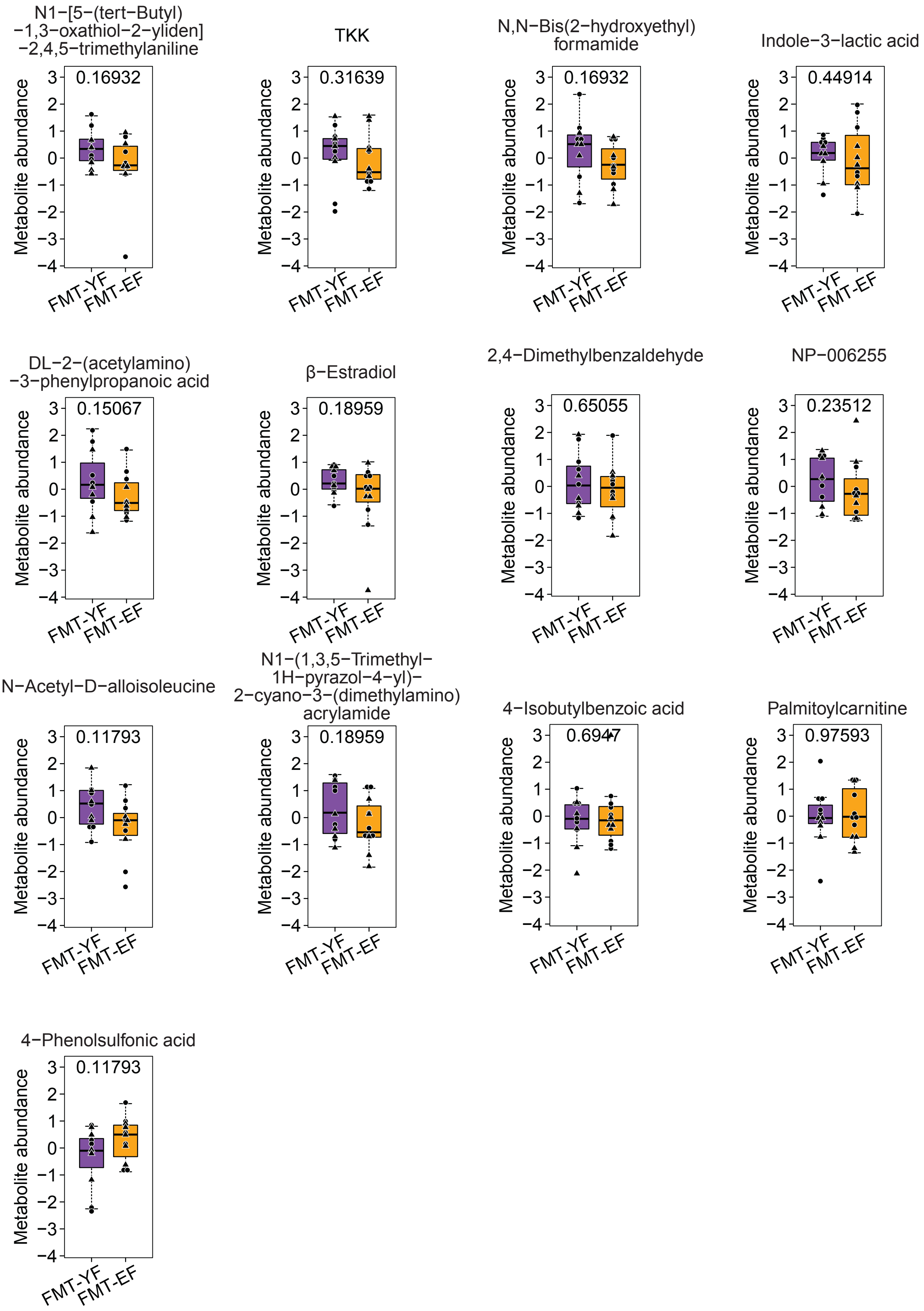

Extended Data Fig. 16

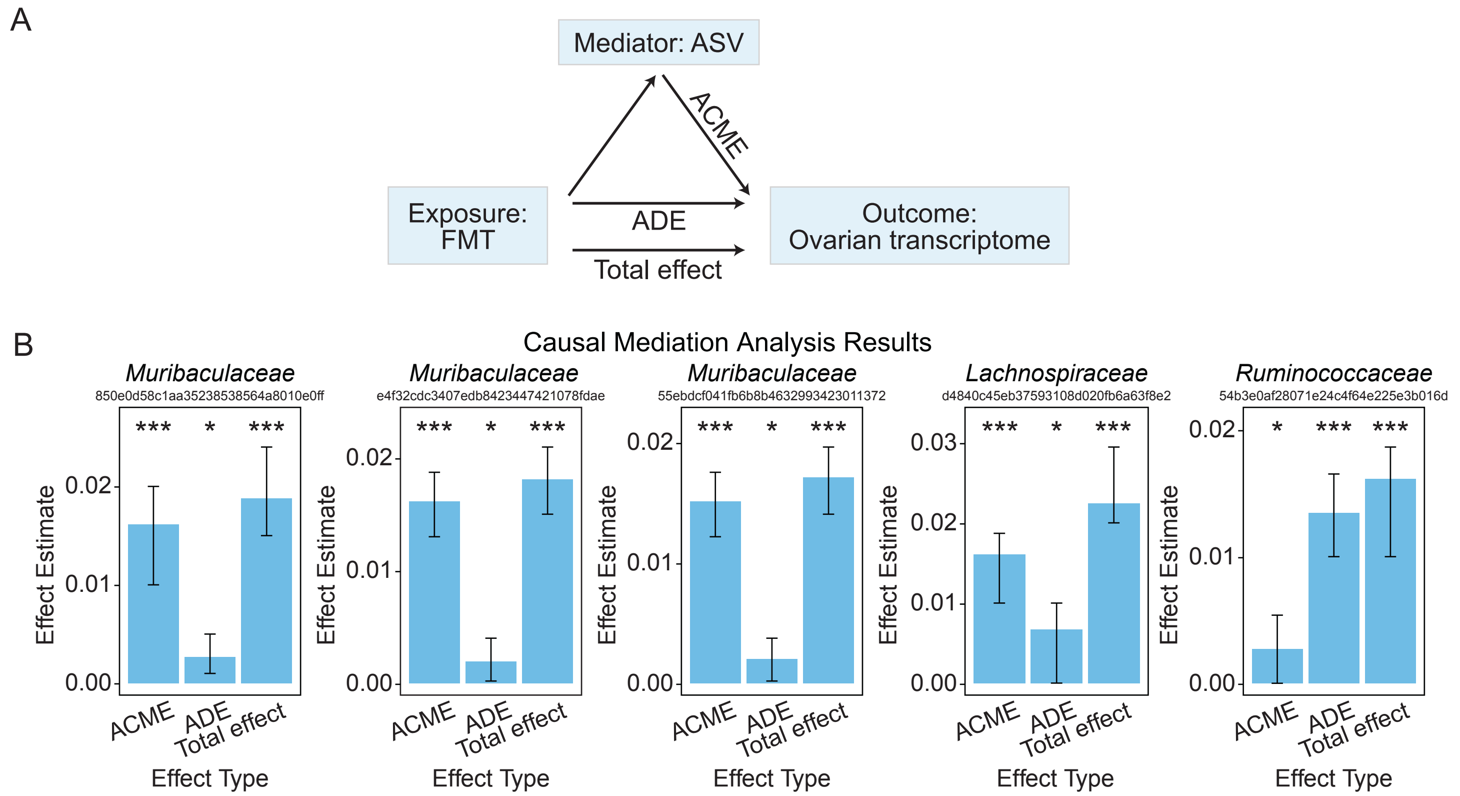
